# The origins and relatedness structure of mixed infections vary with local prevalence of *P. falciparum* malaria

**DOI:** 10.1101/387266

**Authors:** Sha Joe Zhu, Jason A. Hendry, Jacob Almagro-Garcia, Richard D. Pearson, Roberto Amato, Alistair Miles, Daniel J. Weiss, Tim C.D. Lucas, Michele Nguyen, Peter W. Gething, Dominic Kwiatkowski, Gil McVean, for the Pf3k Project

## Abstract

Individual malaria infections can carry multiple strains of *Plasmodium falciparum* with varying levels of relatedness. Yet, how local epidemiology affects the properties of such mixed infections remains unclear. Here, we develop an enhanced method for strain deconvolution from genome sequencing data, which estimates the number of strains, their proportions, identity-by-descent (IBD) profiles and individual haplotypes. Applying it to the Pf3k data set, we find that the rate of mixed infection varies from 29% to 63% across countries and that 51% of mixed infections involve more than two strains. Furthermore, we estimate that 47% of symptomatic dual infections contain sibling strains likely to have been co-transmitted from a single mosquito, and find evidence of mixed infections propagated over successive infection cycles. Finally, leveraging data from the Malaria Atlas Project, we find that prevalence correlates within Africa, but not Asia, with both the rate of mixed infection and the level of IBD.

## 1 Introduction

Individuals infected with malaria-causing parasites of the genus *Plasmodium* often carry multiple, distinct strains of the same species (Bell et al., 2006). Such mixed infections, also known as complex infections, are likely indicative of intense local exposure rates, being common in regions of Africa with high rates of prevalence (Howes et al., 2016). However, they have also been documented for *P. vivax* and other malaria-causing parasites (Ivo Mueller, 2007; Collins, 2012), even in regions of much lower prevalence (Howes et al., 2016; Steenkeste et al., 2010). Mixed infections have been associated with increased disease severity (de Roode et al., 2005) and also facilitate the generation of genomic diversity within the parasite, enabling co-transmission to the mosquito vector where sexual recombination occurs (Mzilahowa et al., 2007). The distribution of mixed infection duration, and whether the clearance of one or more strains results purely from host immunity (Borrmann and Matuschewski, 2011) or can be influenced by interactions between the distinct strains (Enosse et al., 2006; Bushman et al., 2016), are all open questions.

Although mixed infections can be studied from genetic barcodes (Galinsky et al., 2015) or single nucleotide polymorphisms (SNPs) (O’Brien et al., 2016), genome sequencing provides a more powerful approach for detecting mixed infections (Chang et al., 2017). Genetic differences between co-existing strains manifest as polymorphic loci in the DNA sequence of the isolate. The higher resolution of sequencing data allows the use of statistical methods for estimating the number of distinct strains, their relative proportions, and genome sequences (Zhu et al., 2018). Although genomic approaches cannot identify individuals infected multiple times by identical strains, and are affected by sequencing errors and problems of incomplete or erroneous reference assemblies, they provide a rich characterisation of within host diversity (Manske et al., 2012; Auburn et al., 2012; Pearson et al., 2016).

Previous research has highlighted that co-existing strains can be highly related (Nair et al., 2014; Trevino et al., 2017). For example, in *P. vivax*, 58% of mixed infections show long stretches of within host homozygosity (Pearson et al., 2016). In addition, Nkhoma et al. (2012) reported an average of 78.7% *P. falciparum* allele sharing in Malawi and 87.6% sharing in Thailand. A mixed infection with related strains can arise through different mechanisms. Firstly, relatedness is created when distinct parasite strains undergo meiosis in a mosquito vector. A mosquito vector can acquire distinct strains by biting a single multiply-infected individual, or multiple infected individuals in close succession. Co-transmission of multiple meiotic progeny produces a mixed infection in a single-bite, containing related strains. Alternatively, relatedness in a mixed infection can result from multiple bites in a parasite population with low genetic diversity, such as is expected during the early stages of an outbreak or following severe population bottlenecks; for instance, those resulting from an intervention (Mouzin et al., 2010; Wong et al., 2017; Daniels et al., 2015). Interestingly, serial co-transmission of a mixed infection is akin to inbreeding, producing strains with relatedness levels well above those of standard siblings.

The rate and relatedness structure of mixed infections are therefore highly relevant for understanding regional epidemiology. However, progress towards utilising this source of information is limited by three problems. Firstly, while strain deconvolution within mixed infections has received substantial attention (Galinsky et al., 2015; O’Brien et al., 2016; Chang et al., 2017; Zhu et al., 2018), currently, no methods perform both deconvolution of strains and estimation of relatedness. Because existing deconvolution methods assume equal relatedness along the genome, differences in relatedness that occur, for example through infection by sibling strains, can lead to errors in the estimation of the number, proportions and sequences of individual strains (Figure 1). Recently, progress has been made in the case of dual-infections with balanced proportions (Henden et al., 2018), but a general solution is lacking. The second problem is that little is known about how the rate and relatedness structure of mixed infections relates to underlying epidemiological parameters. Informally, mixed infections will occur when prevalence is high; an observation exploited by Cerqueira et al. (2017) when estimating changes in transmission over time. However, the quantitative nature of this relationship, the key parameters that influence mixed infection rates and how patterns of relatedness relate to infection dynamics are largely unexplored. Finally, an important issue, though not one addressed here, is the sampling design. Malaria parasites may be taken from individuals presenting with disease or as part of a surveillance programme. They are also often highly clustered in time and space. What impact different sampling approaches have on observed genomic variation is not clear. Nevertheless, because mixed infection rates are likely to respond rapidly to changes in prevalence (Volkman et al., 2012), exploring these challenges may render critical for malaria control in the field.

**Figure 1:**
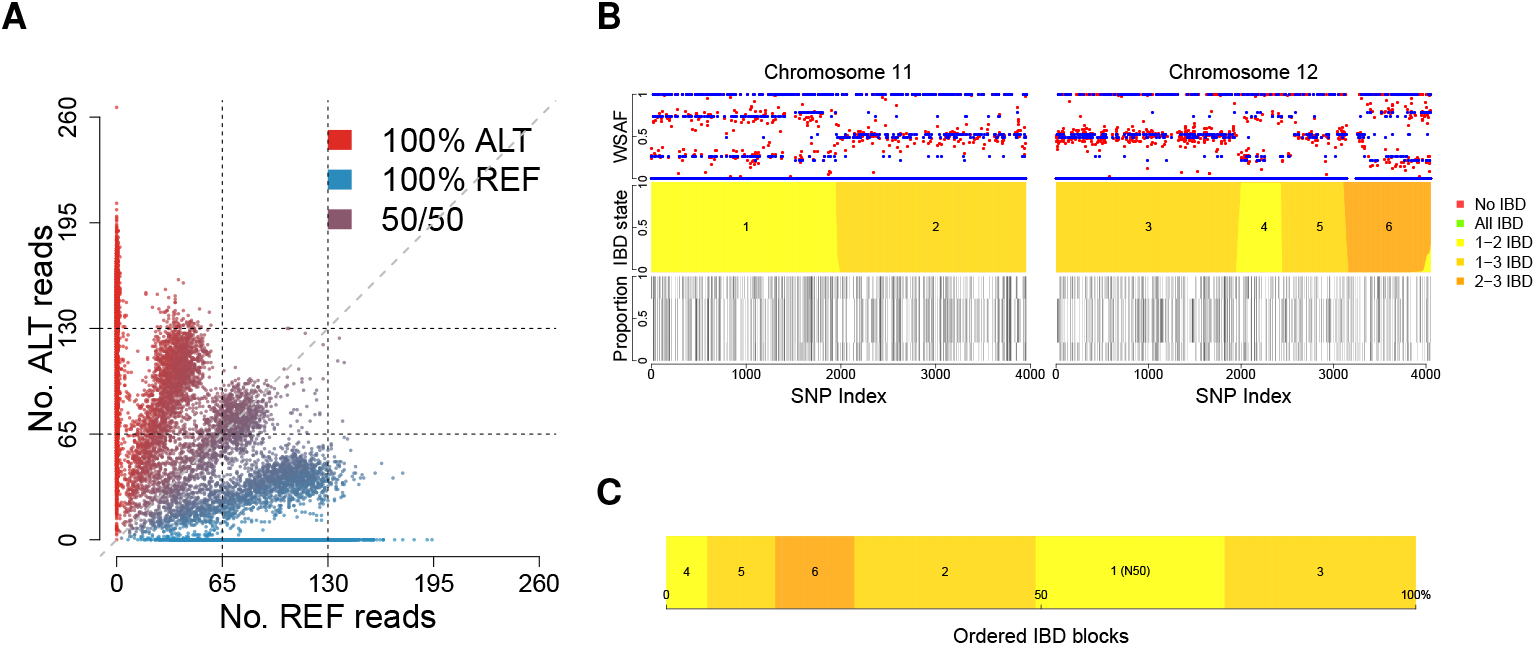
Deconvolution of a complex field sample PD0577-C from Thailand. (A) Scatter-plot showing the number of reads supporting the reference (REF: x-axis) and alternative (ALT: y-axis) alleles. The multiple clusters indicate the presence of multiple strains, but cannot distinguish the exact number or proportions. (B) The profile of within-sample allele frequency along chromosomes 11 and 12 (red points) suggests a changing profile of IBD with three distinct strains, estimated to be with proportions of 22%, 52% and 26% respectively (other chromosomes omitted for clarity, see Figure 1-Supplement ??); blue points indicate expected allele frequencies within the isolate. However, the strains are inferred to be siblings of each other: green segments indicate where all three strains are IBD; yellow, orange and dark orange segments indicate the regions where one pair of strains are IBD but the others are not. In no region are all three strains inferred to be distinct. (C) Statistics of IBD tract length, in particular illustrating the N50 segment length. A graphical description of the modules and workows for `DEploidIBD` is given in Figure 1-Supplement ??.

Here, we develop, test and apply an enhanced method for strain deconvolution called `DEploidIBD`, which builds on our previously-published `DEploid` software. The method separates estimation of strain number, proportions, and relatedness (specifically the identity-by-descent, or IBD, profile along the genome) from the problem of inferring genome sequences. This strategy provides substantial improvements to accuracy when strains are closely related. We apply `DEploidIBD` to 2,344 field isolates of *P. falciparum* collected from 13 countries over a range of years (2001-2014) and available through the Pf3k Project (see Supplementary Note), and characterise the rate and relatedness patterns of mixed infections. In addition, we develop a statistical framework for characterising the processes underlying mixed infections, estimating that nearly half of symptomatic mixed infections arise from the transmission of sibling strains, as well as demonstrating the propagation of mixed infections through multiple cycles of host-vector transmission. Finally, we investigate the relationships between statistics of mixed infection and epidemiological estimates of pathogen prevalence (MAP, 2017), showing that, at a global level, regional rates of mixed infection and levels of background IBD are correlated with estimates of malaria parasite prevalence.

## 2 Strain deconvolution in the presence of relatedness

Existing methods for deconvolution of mixed infections typically assume that the different genetic strains present in mixed infections are unrelated. This assumption allows for efficient computation of priors for allele frequencies within samples, either through assuming independence of loci (O’Brien et al., 2016) or as sequences generated as imperfect mosaics of some (predefined) reference panel (Zhu et al., 2018). However, when strains are related to each other, and particularly when patterns of IBD vary along the genome (for example through being siblings), the constraints imposed on within-sample allele frequencies through IBD can cause problems for deconvolution methods, which can try to fit complex strain combinations (with relatedness) as simpler configurations (without relatedness). Below we outline the approach we take to integrating IBD into `DEploid`. Further details are provided in the Supplementary Materials.

### 2.1 Decoding genomic relatedness among strains

A common approach to detecting IBD between two genomes is to employ a hidden Markov Model that transitions into and out of IBD states (Chang et al., 2015; Gusev et al., 2009, 2011). We have generalised this approach to the case of *K* haploid *Plasmodium* genomes (strains). In this setting, there are 2^*K*^ possible genotype configurations, as each of the *K* strains can be either reference (i.e. same as the reference genome used during assembly), or alternative (i.e. carry a different allele) at a given locus (we assume all variation is bi-allelic). If each of the *K* strains constitutes a unique proportion of the infection, each genotype configuration will produce a distinct alternative within sample allele frequency (WSAF; Figure 1A), which defines the expected fraction of total sequencing reads that are alternative at a given locus in the sequenced infection.

The effect of IBD among these *K* strains is to limit the number of distinct genotype configurations possible, in a way that depends on the pattern of IBD sharing. Consider that, for any given locus, the *K* strains in the infection are assigned to *j ≤ K* possible reference haplotypes. IBD exists when two or more strains are assigned to the same haplotype. In this scenario, the total number of possible patterns of IBD for a given *K* is equal to 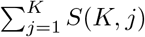, where *S*(*K, j*) is the number of ways *k* objects can be split into *j* subsets; a Stirling number of the second kind (Graham et al., 1988). Thus, for two strains, there are two possible IBD states (IBD or non-IBD), for three strains there are five states (all IBD, none IBD and the three pairwise IBD configurations), for four strains there are fifteen states (see Supplementary Materials), and so on. We limit analysis to a maximum of four strains for computational efficiency. Finally, for a given IBD state, only 2^*j*^ rather than 2^*K*^ genotype configurations are possible, thereby restricting the set of possible WSAF values.

Moving along the genome, recombination can result in changes in IBD state, hence changing the set of possible WSAF values at loci (Figure 1B). To infer IBD states we use a hidden Markov model, which assumes linkage equilibrium between variants for computational efficiency, with a Gamma-Poisson emission model for read counts to account for over-dispersion (see Supplementary Materials). Population-level allele frequencies are estimated from isolates obtained from a similar geographic region. Given the structure of the hidden Markov model, we can compute the likelihood of the strain proportions by integrating over all possible IBD sharing patterns, yielding a Bayesian estimate for the number and proportions of strains (see Supplementary material S1.4 Implementation details). We then use posterior decoding to infer the relatedness structure across the genome (Figure 1B). To quantify relatedness, we compute the mean IBD between pairs of strains and statistics of IBD tract length (mean, median and N50, the length-weighted median IBD tract length, Figure 1C).

In contrast to our previous work, `DEploidIBD` infers strain structure in two steps. In the first we estimate the number and proportions of strains using Markov Chain Monte-Carlo (MCMC), allowing for IBD as described above. In the second, we infer the individual genomes of the strains, using the MCMC methodology of Zhu et al. (2018), which can account for linkage disequilibrium (LD) between variants, but without updating strain proportions. The choice of reference samples for deconvolution is described in Zhu et al. (2018) and in the Supplementary Materials. During this step we do not use the inferred IBD constraints *perse*, though the inferred haplotypes will typically copy from the same (or identical) members of the reference panel within the IBD tract.

## 3 Results

### 3.1 Method validation

#### 3.1.1 Validation using experimentally generated mixed infections

We first sought to characterise the behaviour of `DEploidIBD` and compare its performance to the previously published method, `DEploid`. To this end, we first re-analysed a set of 27 experimentally generated mixed infections (Wendler, 2015) that had been previously deconvoluted by `DEploid` (Zhu et al., 2018) using `DEploidIBD` (Figure 2-supplement **??**). These mixed infections were created with combinations of two or three laboratory strains (selected from 3D7, Dd2, HB3 and 7G8), set at varying known proportions (Wendler, 2015), and therefore provide a simple framework for evaluating inference of the number of strains (*K*) and their proportions. Since the method allows deconvolution of mixed infections containing up to four strains, we augmented the experimental mixtures by combining all four lab strains *in silico* at differing proportions (see Supplementary material S2.1 *In silico* lab mixtures). Using this approach, we found that `DEploid` and `DEploidIBD` performed comparably, except in the case of three strains with equal proportions, where LD information is necessary to achieve accurate deconvolution and `DEploid` performed better. Both `DEploid` and `DEploidIBD` struggled to deconvolute our *in silico* mixtures of four strains, typically underestimating the number of strains present.

**Figure 2:**
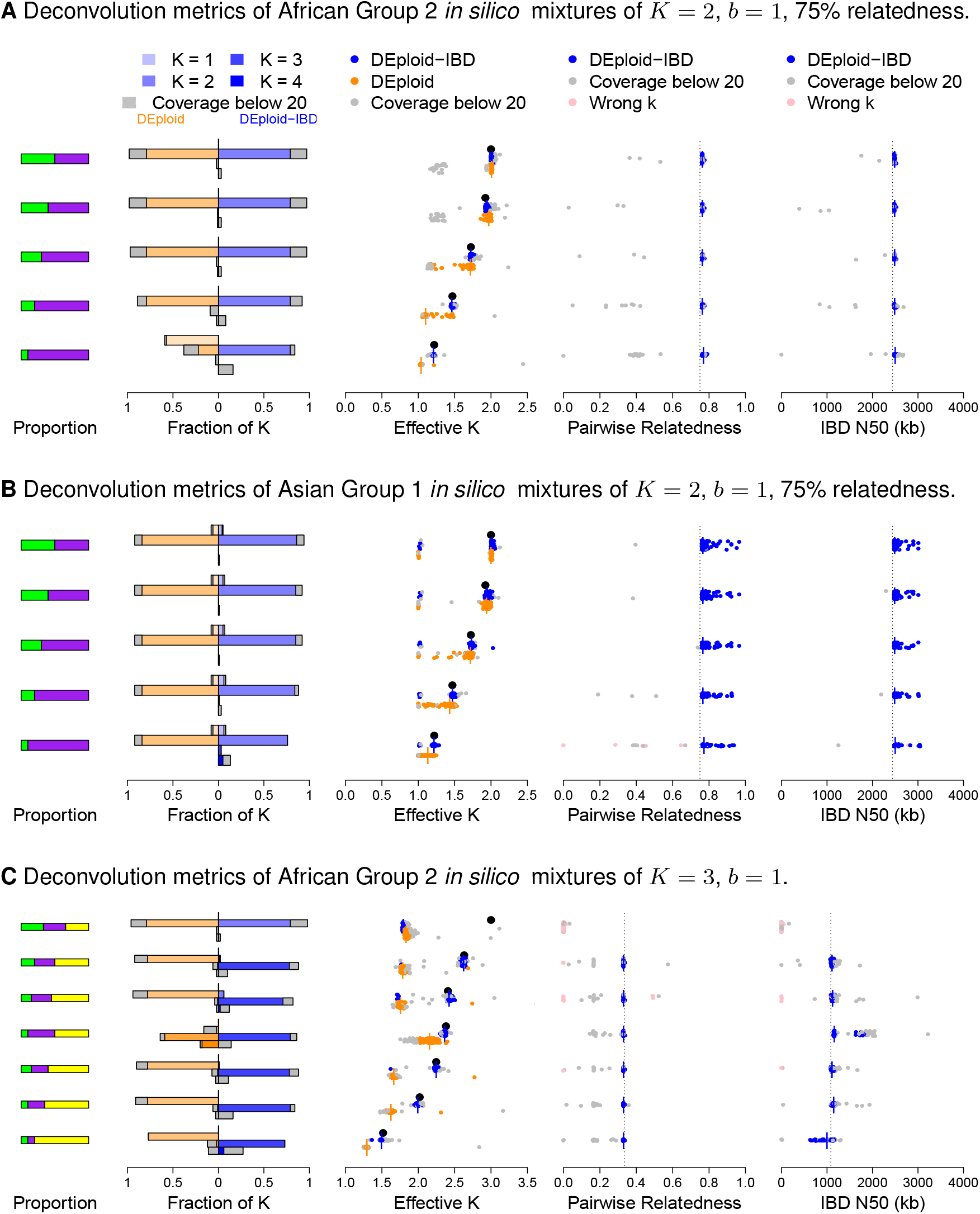
Performance of `DEploidIBD` and `DEploid` on 100 *in silico* mixtures for each of three different scenarios. From the left to the right, the panels show the strain proportion compositions, distribution of inferred *K* using both methods: `DEploid` in orange and `DEploidIBD` in blue, effective number of strains, pairwise relatedness and IBD N50 (the latter two only for `DEploidIBD`). From top to the bottom, cases are ordered from even strain proportions to the most imbalanced composition. Grey points identify experiments of low coverage data, and pink identify cases where *K* is inferred incorrectly. (A) *In silico* mixtures of two African strains with high-relatedness (75%) for 7,757 (s.d. 178) sites on Chromosome 14, Note that `DEploid` underestimates the minor strain proportion if strains have high relatedness. In the extreme case, `DEploid` misclassifies a *K* = 2-mixture as clonal, whereas `DEploidIBD` consistently estimates the correct proportions. (B) *In silico* mixtures of two Asian strains with high-relatedness (75%) for 3,041 sites (s.d. 227) on Chromosome 14, Note that `DEploid` underestimates strain number when the minor strain is low frequency, while `DEploid` typically performs well. (C) *In silico* mixtures of three African strains, where each pair is IBD over a distinct third of the chromosome. Note that both methods fail to deconvolute the case of equal proportions. However, for unbalanced mixtures, `DEploidIBD` consistently performs better than `DEploid`.

#### 3.1.2 Validation against simulated mixed infections

Validation using mixtures of lab strains has two limitations: (i) the strains comprising the mixed infection were part of the reference panel and (ii) no IBD was present. We therefore investigated the ability of `DEploidIBD` to recover IBD between strains within a mixed infection, in the context of a realistic reference panel, and with strains typical of those we observe in nature. To achieve this, we designed a validation framework where clonal samples from the Pf3k project were combined *in silico* to produce simulated mixed infections, allowing us to create examples with varying numbers of strains and proportions, and to introduce tracts of IBD, by copying selected sections of the genome between strains. Using this framework, we constructed a broad suite of simulated mixed infections, derived from clonal samples from Africa and Asia that were combined into mixtures of 2, 3 and 4 strains with variable proportions and IBD configurations.

We randomly selected 189 clonal samples of African origin and 204 clonal samples of Asian origin from which to construct our simulated mixed infections and restricted the analysis to chromosome 14 to reduce computational time. Starting with mixed infections containing two strains, we randomly took two samples of African or Asian origin and combined them at proportions ranging from highly imbalanced (10% and 90%) to exactly balanced (each 50%) and used copying to produce either no (0%), low (25%), medium (50%) or high (75%) levels of IBD (note that background IBD between the two clonal strains may also exist). In total this resulted in 4,000 *K* = 2 mixed infections, each of which was deconvoluted with `DEploid` and `DEploidIBD`. Outputs of `DEploidIBD` were compared to the true values for each simulated infection, including the inference of *K*, the effective *K*_*e*_ (computed as 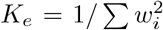, where *w*_*i*_ is the proportion of the *i*th strain, thus incorporating proportion inference), the average pairwise relatedness between strains (for *K* = 2, this is the fraction of the genome inferred to be IBD), and the inference of IBD tract length, expressed as the IBD N50 metric.

For mixtures of two strains, both `DEploid` and `DEploidIBD` performed well in scenarios where the IBD between strains was low (¡=25%). In moderate or high IBD scenarios with imbalanced strain proportions, `DEploid` tended to underestimate the proportion of the minor strain resulting in underestimation of *K*_*e*_, whereas `DEploidIBD` was able to infer the proportion of these mixtures correctly (Figure 2). The main novelty of `DEploidIBD` is the calculation of an IBD profile between strains. We found that the IBD summary statistics produced by `DEploidIBD` were accurate across all two-strain mixed infections tested in Africa. In Asia, `DEploidIBD` tended to estimate more IBD than was simulated (Figure 2B). However, this likely reflects the presence of higher background IBD in Asia rather than systematic error.

To simulate realistic mixed infections containing 3 or 4 strains, we first considered the different transmission scenarios under which they can arise. We modelled a mixed infection of *K* strains as resulting from *b* biting events, where *K ∈* {3, 4} and 1 ≤ *b* ≤ *K*. When greater than one strain is transmitted in a single biting event, the co-transmitted strains will share IBD, as a consequence of meiosis occurring in the mosquito. Strains transmitted through independent bites, causing superinfection in the host, do not share any IBD beyond background. Following this paradigm, we generated a suite of mixed infection types: *K* = 3, *b* = 1, 2, 3 and *K* = 4, *b* = 1, 2, 2, 3, 4 (the first *b* = 2 has two strains per bite, the second three and one); and simulated each of these across a variety of proportions, again using sets of clonal samples from Africa and Asia as starting strains.

As with the experimental validation, the balanced-proportion *K* = 3 mixed infections generated *in silico* proved challenging to deconvolute, with both methods inferring the presence of two rather than three strains (Figure 2C). In mixed infections with imbalanced proportions, we found that, in African samples with IBD (*b* = 1, 2), `DEploid` tended to either underestimate the number of strains present, or infer proportions incorrectly. In Asian samples this is less of an issue as the reference panels can provide better prior information for deconvolution due to lower diversity. In contrast, `DEploidIBD` consistently gave the correct number of strains and proportions in such cases, and produced IBD statistics that were accurate as long as the median coverage of simulated infections was > 20x. Both methods struggled to deconvolute mixed infections of four strains (Figure 2-supplement **??**), although performed better (i.e. inferred *K* = 4 greater than 50% of the time) for mixtures with less IBD (*b* = 3, 4). However, even in these cases, estimates of the proportions and IBD statistics were variable, indicating that further work is needed before *K* = 4 mixed infections can be reliably deconvoluted.

Finally, we used the *in silico* approach to explore the quality of haplotypes inferred by `DEploidIBD`, focusing on *K* = 2 infections across variable proportions. We compared the haplotype inferences between `DEploid` and DEpoloidIBD using the error model described in the Supplementary Material, and found that rates of genotype error are similar for the two approaches in settings of low relatedness (`DEploidIBD` has an error rate of 0.7% per site per strain for 20/80 mixtures and 1.4% for 50/50 mixtures). However, for the 20/80% mixtures with high relatedness, genotype error for `DEploid` increased to 1.8%, while remaining at 0.8% for `DEploidIBD` (Figure 3A). Switch errors in haplotype estimation are comparable between the two methods and decrease with increasing relatedness due to higher homozygosity (Figure 3B). Finally, we identified a simple metric to compute on inferred haplotypes that can identify low quality haplotypes (see Supplementary Material),

**Figure 3:**
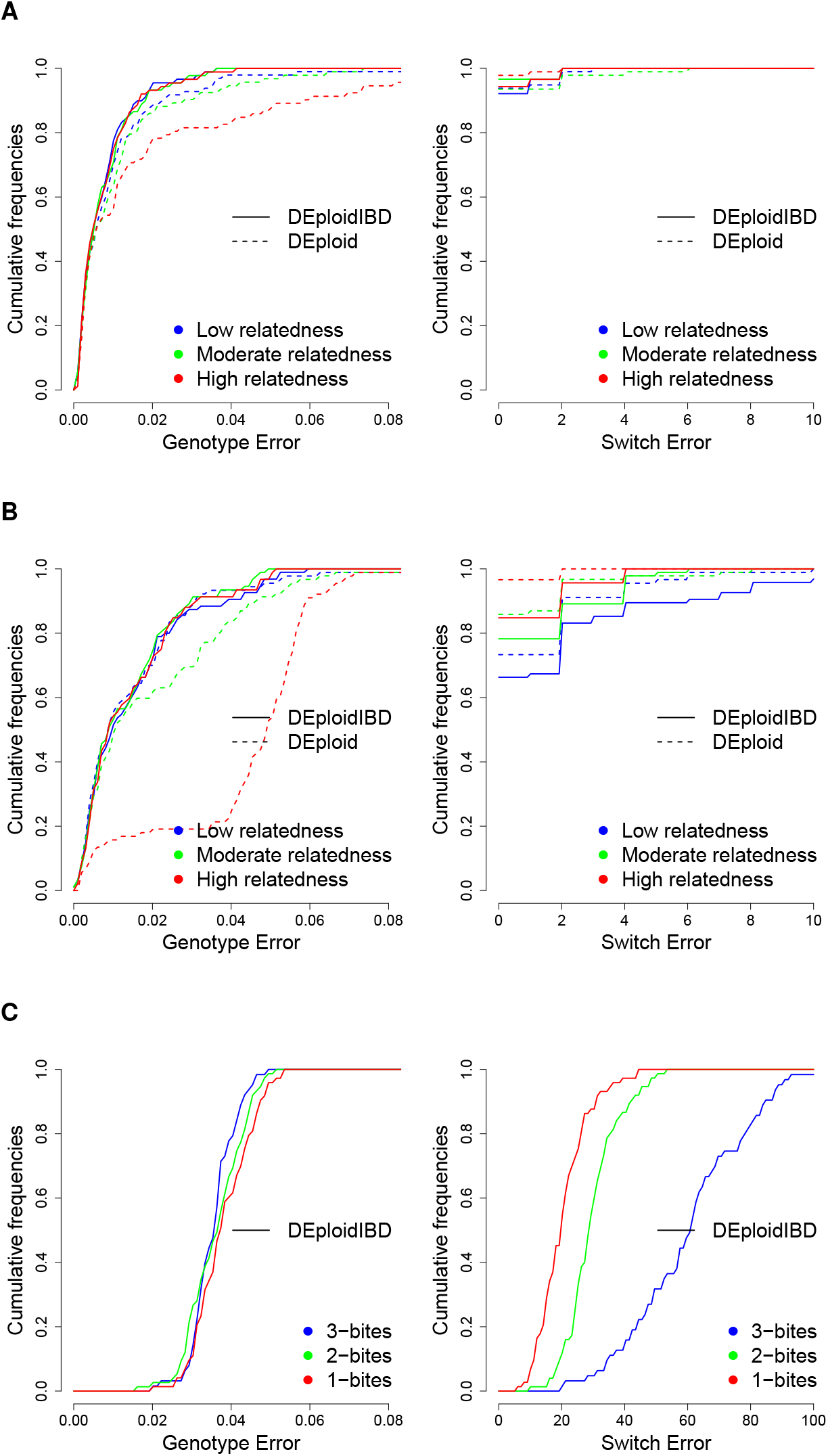
Cumulative distribution of the average per site genotype error (left) and switch error (right) across simulated mixtures. (A) Error rates of Asian *in silico* samples of three levels of IBD (25%, 50% and 75%) for a *K* = 2 mixture with proportions of 20*/*80%. Because `DEploidIBD` estimates proportions more accurately, it enables better haplotype inference. (B) Error rates of African *in silico* samples of three levels of IBD (25%, 50% and 75%) for a *K* = 2 mixture with proportions of 20*/*80%. Inference in Asia benefits from better reference panels (due to lower overall diversity) and therefore gives lower error rates than in Africa. (C) `DEploidIBD` error rates for African *in silico* samples of three mosquito biting scenarios for a *K* = 3 mixture with proportions of 10*/*10*/*80%. The additional strain increases the difficulty of haplotype inference, particularly in the case of three independent bites.

### 3.2 Geographical variation in mixed infection rates and relatedness

To investigate how the rate and relatedness structure of mixed infections varies among geographical regions with different epidemiological characteristics, we applied `DEploidIBD` to 2,344 field samples of *P. falciparum* released by the Pf3k project (Pf3k Consortium, 2016). These samples were collected under a wide range of studies with differing designs, though the majority of samples were collected from symptomatic individuals seeking clinical treatment. An important exception are the samples from Senegal which, though collected passively at a clinic, were screened to contain only one strain by SNP barcode (Daniels et al., 2015). A summary of the data sources is presented in Table 1 and full details regarding study designs can be found at https://www.malariagen.net/projects/pf3k#sampling-locations. Details of data processing are given in the Supplementary Material. For deconvolution, samples were grouped into geographical regions by genetic similarity; four in Africa, and three in Asia. (Table 1). Reference panels were constructed from the clonal samples found in each region. Since previous research has uncovered strong population structure in Cambodia (Miotto et al., 2013), we stratified samples into West and North Cambodia when performing analysis at the country level. Diagnostic plots for the deconvolution of all samples can be found at https://github.com/mcveanlab/mixedIBD-Supplement and inferred haplotypes can be accessed at ftp://ngs.sanger.ac.uk/production/pf3k/technical_working/release_5/mixedIBD_paper_haplotypes/. We identified 787 samples where low sequencing coverage or the presence of low-frequency strains resulted in unusual haplotypes (see Supplementary Material). Estimates of strain number, proportions and IBD states from these samples are used in subsequent analyses, but not the haplotypes. We also confirmed that reported results are not affected by the exclusion of samples with haplotypes with low confidence (data not shown). In all following analyses, only strains present with a proportion of ¿1% in a sample are reported.

**Table 1:**
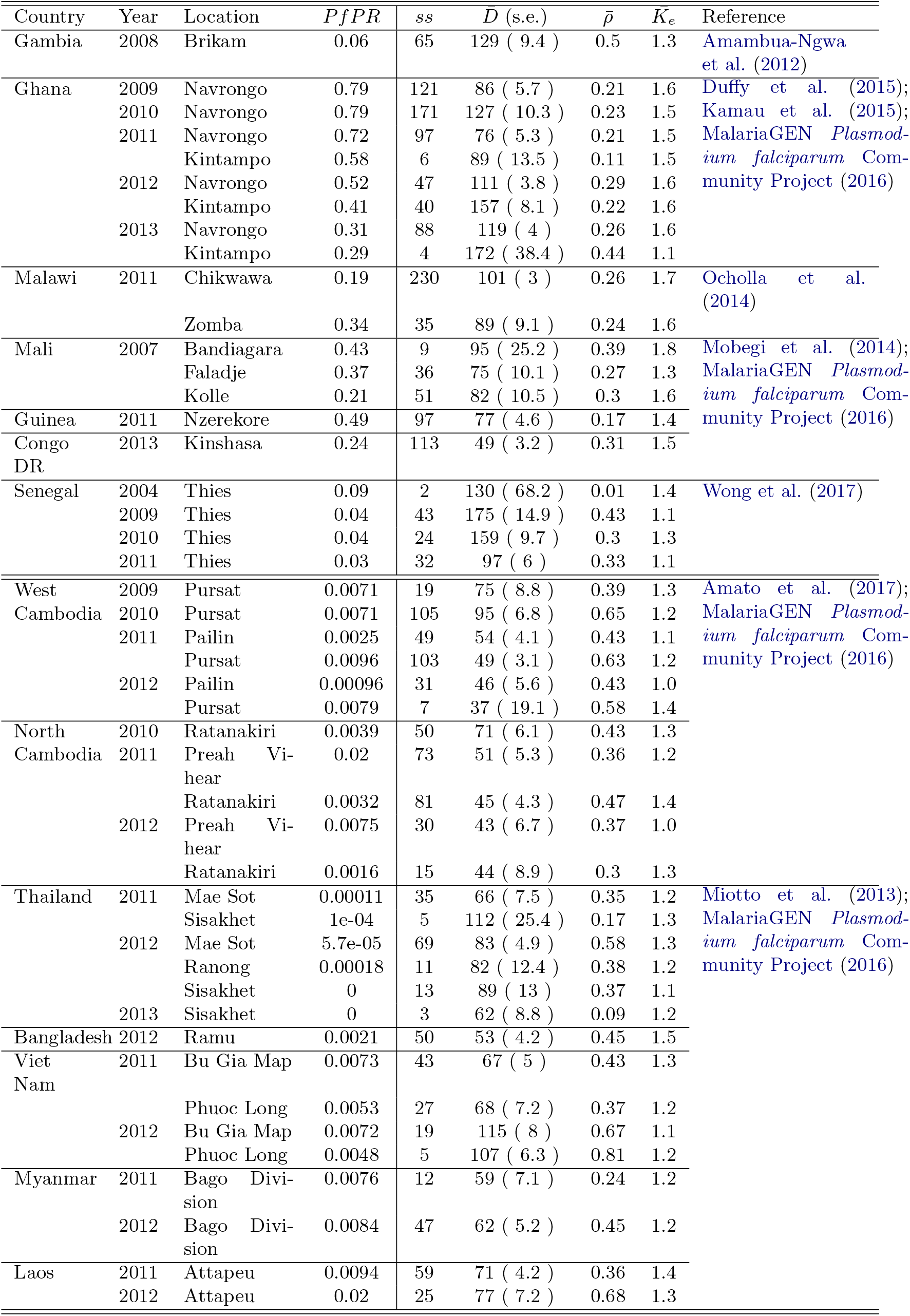
Summary of Pf3k samples. Table 2: Summary of Pf3k samples in data release 5.1, where *D* denotes mean read depth and *ss* is sample size. Genotyping, including both indel and SNP variants, was performed using a pipeline based on GATK best practices, see Methods. Data available from ftp://ngs.sanger.ac.uk/production/pf3k/release_5/5.1. *Pf PR* is the inferred parasite prevalence rate in a 5 × 5 km resolution grid from the MAP project, centred at the Pf3k sample collection sites; Relatedness *ρ* and effective number of strains *K*_*e*_ are summary metrics from `DEploidIBD` output.

We find substantial variation in the rate and relatedness structure of mixed infections across continents and countries. Within Africa, rates of mixed infection vary from 29% in The Gambia to 63% in Malawi (Figure 4A). Senegal has a rate of mixed infection (18%) lower than The Gambia, however as these samples were screened by SNP barcode to be clonal, this rate should be an underestimate. In Southeast Asian samples, mixed infection rates are in general lower, though also vary considerably; from 21% in Thailand to 54% in Bangladesh. Where data for a location is available over multiple years, we find no evidence for significant fluctuation over time (though we note that these studies are typically not well powered to see temporal variations). We observe that between 5.1% (Senegal) and 40% (Malawi) of individuals have infections carrying more than two strains.

**Figure 4:**
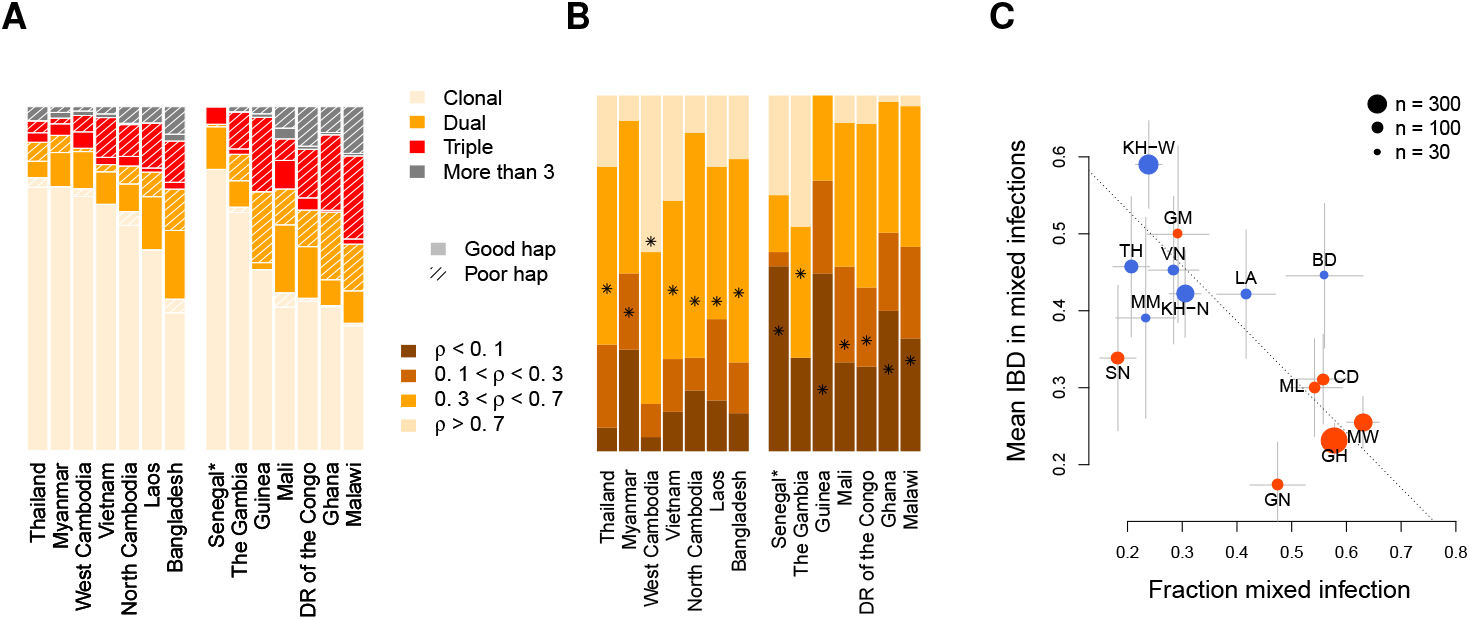
Characterisation of mixed infections across 2,344 field samples of *Plasmodium falciparum*. (A) The fraction of samples, by population, inferred by `DEploidIBD` to be *K* = 1 (clonal), *K* = 2 (dual), *K* = 3 (triple), or *K* = 4 (More than 3). Populations are ordered by rate of mixed infections within each continent. We use shaded regions to indicate the distribution of 787 samples that have low-confidence deconvoluted haplotypes. Senegal is marked with an asterisks as these samples were screened to be clonal. (B) The distribution of average pairwise IBD sharing within mixed infections (including dual, triple and quad infections), broken down into unrelated (where the fraction of the genome inferred to be IBD, *ρ*, is < 0.1), low IBD (0.1 ≥ *ρ* < 0.3), sib-level (0.3 ≥ *ρ* < 0.7) and high (*ρ* ≥ 0.7). Stars indicate the average IBD scaled between 0 and 1 from bottom to the top. Populations follow the same order as in Panel A. (C) The relationship between the rate of mixed infection and level of IBD. Populations are coloured by continent, with size reflecting sample size and error bars showing ±1 s.e.m. The dotted line shows the slope of the regression from a linear model. Abbreviations: SN-Senegal, GM-The Gambia, NG-Nigeria, GN-Guinea, CD-The Democratic Republic of Congo, ML-Mali, GH-Ghana, MW-Malawi, MM-Myanmar, TH-Thailand, VN-Vietnam, KH-Cambodia, LA-Laos, BD-Bangladesh.

Relatedness between samples and populations also varies substantially. In dual infections, the average fraction of the genome inferred to be IBD ranges from 14% in Guinea to 65% in West Cambodia (Figure 4B). Asian populations show, on average, a higher level of relatedness within dual infections (44%) compared to African populations (26%). Levels of IBD in samples with three or more strains are comparable to those seen in dual infections (average IBD being 45% in Asia and 37% in Africa) and significantly correlated at the country level, with correlation of 0.75(P = 0.0019, weighted by the number of mixed samples). Overall, 51% of all mixed infections involve strains with over 30% of the genome being IBD.

We next considered the relationship between mixed infection rate and the level of IBD. We find that populations with higher rates of mixed infection tend to have lower levels of IBD within mixed infections (linear model P = 0.06 after accounting for a continental level difference and weighted by sample size). However, the continental level effect is driven by Senegal, which has an unusual combination of low mixed infections and also low IBD. Excluding Senegal, we find a consistent pattern across populations (Figure 4C), with a strong negative correlation between mixed infection rate and the level of IBD (Pearson *r* = −0.65, P = 3 × 10^−4^). Previous work has demonstrated how a recent and dramatic decline in *P. falciparum* prevalence within Senegal has left an impact on patterns of genetic variation (Daniels et al., 2015), which may explain its unusual profile.

### 3.3 Inferring the origin of IBD in mixed infections

The high levels of IBD observed in many mixed infections suggest the presence of sibling strains (Figure 5). To quantify the expected IBD patterns between siblings, we developed a meiosis simulator for *P. falciparum* (`pf-meiosis`), incorporating relevant features of malaria biology that can impact the way IBD is produced in a mosquito and detected in a human host. Most importantly, a single infected mosquito can undergo multiple meioses in parallel, one occurring for each oocyst that forms on the mosquito midgut (Ghosh et al., 2000). In a mosquito infected with two distinct strains, each oocyst can either self (the maternal and paternal strain are the same) or outbreed (the maternal and paternal strains are different). We model a *K* = *n* mixed infection as a sample of *n* strains (without replacement, as drawing identical strains yields *K* = *n* − 1) from the pool of strains created by all oocysts. Studies of wild-caught *Anopheles Gambiae* suggest that the distribution of oocysts is roughly geometric, with the majority of infected mosquitoes carrying only one oocyst (Beier et al., 1991; Collins et al., 1984). In such a case, we find that a *K* = 2 infection will have an expected IBD of 1*/*3, consistent with the observations of Wong et al. (2018). Conditioning on at least one progeny originating from an outbred oocyst (such that a detectable recombination event has occurred), the expected IBD asymptotically approaches 1*/*2 as the total number of oocysts grows (see Supplementary Materials).

**Figure 5:**
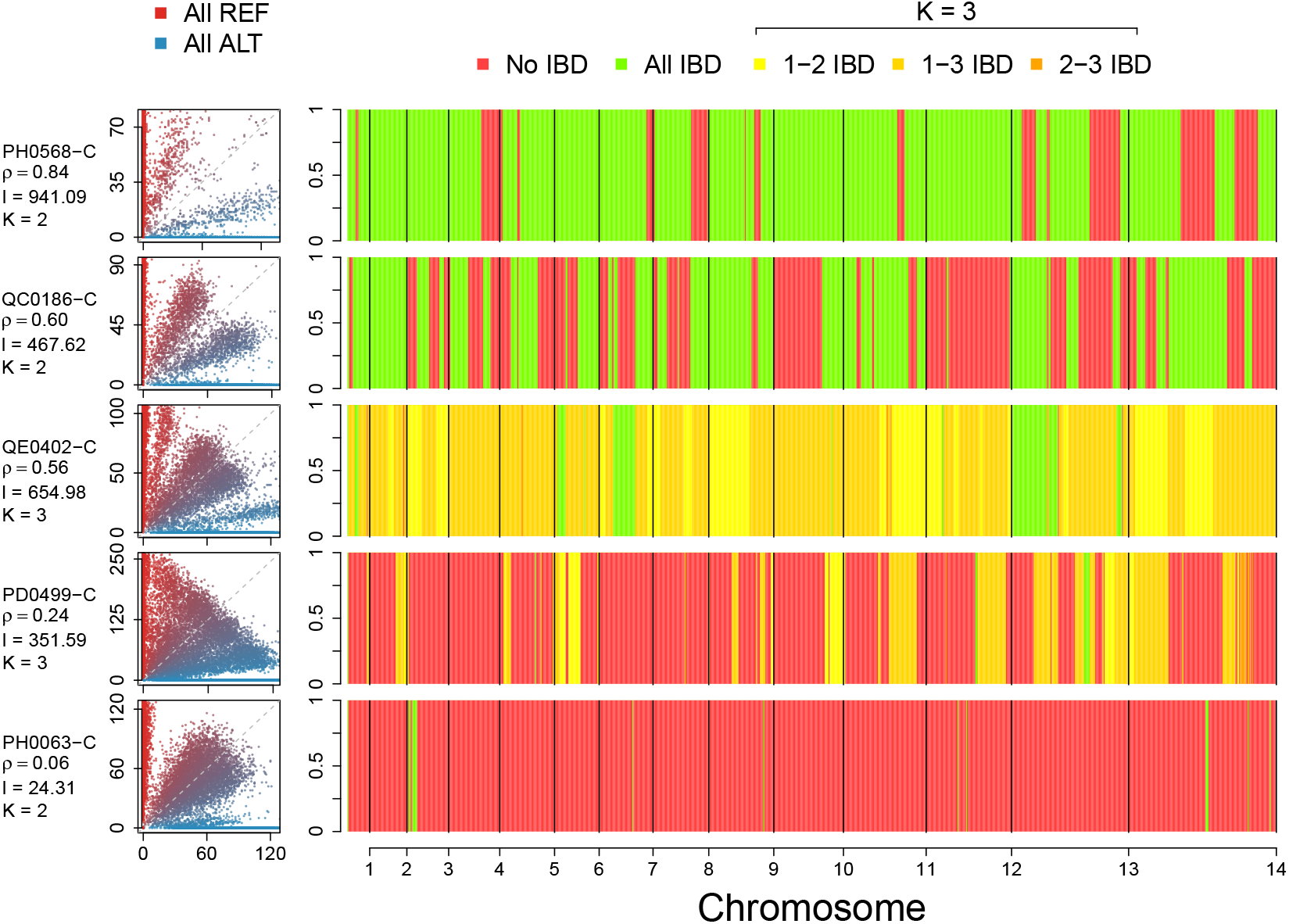
Example IBD profiles in mixed infections. Plots showing the ALT versus REF plots (left hand side) and inferred IBD profiles along the genome for five strains of differing composition. From top to bottom: A dual infection of highly related strains (*ρ* = 0.84); a dual infection of two sibling strains (*ρ* = 0.6); a triple infection of three sibling strains (note the absence of stretches without IBD); a triple infection of two related strains and one unrelated strain; and a triple infection of three unrelated strains. The numbers below the sample IDs indicate the average pairwise IBD, *r*, the mean length of IBD segments, *l*, in kb and the inferred number of distinct strains, *K*, respectively.

Using this simulation framework, we sought to classify observed mixed infections based on their patterns of IBD. We used two summary statistics to perform the classification: mean IBD segment length and IBD fraction. We built empirical distributions for these two statistics for each country in Pf3k, by simulating meiosis between pairs of clonal samples from that country. In this way, we control for variation in genetic diversity (as background IBD between clonal samples) in each country. Starting from a pair of clonal samples (*M* = 0, where *M* indicates the number of meioses that have occurred), we simulated three successive rounds of meiosis (*M* = 1, 2, 3), representing the creation and serial transmission of a mixed infection (Figure 6A). Each round of meiosis increases the amount of observed IBD. For example, in Ghana, the mean IBD fraction for *M* = 0 was 0.002, for *M* = 1 was 0.41, for *M* = 2 was 0.66, and for *M* = 3 was 0.80 (Figure 6B). West Cambodia, which has lower genetic diversity, had a mean IBD fraction of 0.08 for *M* = 0 and consequently, the mean IBD fractions for higher values of *M* were slightly increased, to 0.46, 0.68, 0.81 for *M* = 1, 2 and 3, respectively (Figure 6B).

**Figure 6:**
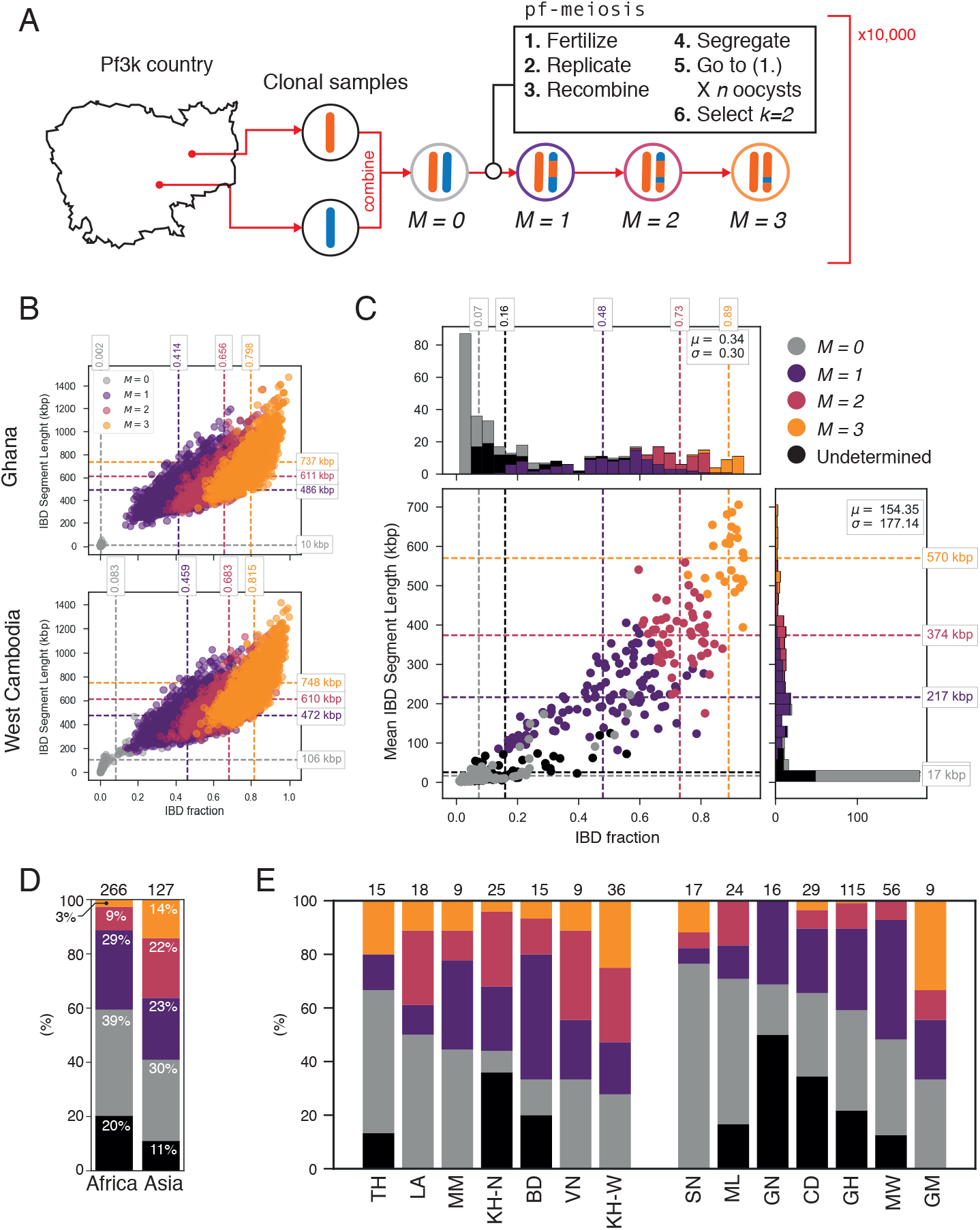
Identifying sibling strains within mixed infections. (A) Schematic showing how IBD fraction and IBD segment length distributions are created for *k* = 2 mixed infections using `pf-meiosis`. Two clonal samples from a given country are combined to create an unrelated (*M* = 0, where *M* is number of meioses that have occurred) mixed infection. The *M* = 0 infection is then passed through 3 rounds of `pf-meioses` to generate *M* = 1, 2, 3 classes, representing serial transmission of the mixed infection (*M* = 1 are siblings). (B) Simulated IBD distributions for *M* = 0, 1, 2, 3 for Ghana (top) and West Cambodia (bottom). A total of 10,000 mixed infections are simulated for each class, from 500 random pairs of clonal samples. (C) Classification results for 393 *K* = 2 mixed infections from 13 countries. Undetermined indicates mixed infections with IBD statistics that were never observed in simulation. (D) Breakdown of class percentage by continent. Total number of samples is given above bars. Colours as in panel C (*M* = 0, grey; *M* = 1, purple; *M* = 2, pink; *M* = 3, orange; Undeteremined, black). (E) Same as (D), but by country. Abbreviations as in Figure 4.

With these simulated distributions, we used Naive Bayes to classify *k* = 2 mixed infections in Pf3k (Figure 6C). Of the 393 *k* = 2 samples containing only high-quality haplotypes (see Supplementary Materials), 325 (83%) had IBD statistics that fell within the range observed across all simulated *M*. Of these, nearly half (183, 47%) were classified as siblings (*M* > 0). Moreover, we observe geographical differences in the rate at which sibling and unrelated mixed infections occur. Notably, in Asia a greater fraction of all mixed infections contained siblings (59% vs. 41% in Africa), driven by a higher frequency of *M* = 2 and *M* = 3 mixed infections (Figure 6D). Mixed infections classified as *M* > 1 are produced by serial co-transmission of parasite strains, i.e. a chain of mixed infections along which IBD increases.

### 3.4 Characteristics of mixed infections correlate with local parasite prevalence

To assess how characteristics of mixed infections relate to local infection intensity, we obtained estimates of *P. falciparum* prevalence (standardised as *Pf PR*_2–10_, prevalence in the 2-to-10 year age range) from the Malaria Atlas Project (MAP, 2017, see Table 1). The country-level prevalence estimates range from 0.01% in Thailand to 55% in Ghana, with African countries having up to two orders of magnitude greater values than Asian ones (mean of 36% in Africa and 0.6% in Asia). However, seasonal and geographic fluctuations in prevalence mean that, conditional on sampling an individual with malaria, local prevalence may be much higher than the longer-term (and more geographically widespread) country-level average, hence we extracted the individual pixel-level estimate of prevalence (corresponding to a 5×5km region) from MAP nearest to each genome collection point. We summarise mixed infection rates by the average effective number of strains, which reflects both the number and proportion of strains present.

Given that samples from Senegal were screened to be primarily single-genome (Daniels et al., 2015), we computed all correlations with prevalence including (*r*_*S*+_) and excluding them (*r*_*S*−_; Figure 7). We find that the effective number of strains is a significant predictor of *Pf PR*_2–10_ globally (*r*_*S*+_ = 0.65, *p* < 10^−5^) and in African populations when Senegal is included (*r*_*S*+_ = 0.48, *p* = 0.04; *r*_*S*−_ = 0.18, *p* = 0.51), but is uncorrelated across Asia. Similarly, *Pf PR*_2–10_ is negatively correlated with background IBD globally (*r*_*S*+_ = −0.43, *p* = 0.004) and across Africa but not in Asia. Surprisingly, the amount of IBD observed within *k* = 2 mixed infections was not correlated with prevalence in Africa or Asia. The rate of sibling infection (*M* = 1) is not correlated with the parasite prevalence (Asia: *r*_*S*+_ = 0.23, *p* = 0.2, Africa: *r*_*S*+_ = 0.16, *p* = 0.5). However, the rate of supersiblings (*k* = 2, *M >* 1) is significantly correlated with *Pf PR*_2–10_ (*r*_*S*+_ = −0.31, *p* = 0.04) at the global scale, suggesting that serial co-transmission may occur more readily in low prevalence regions.

**Figure 7:**
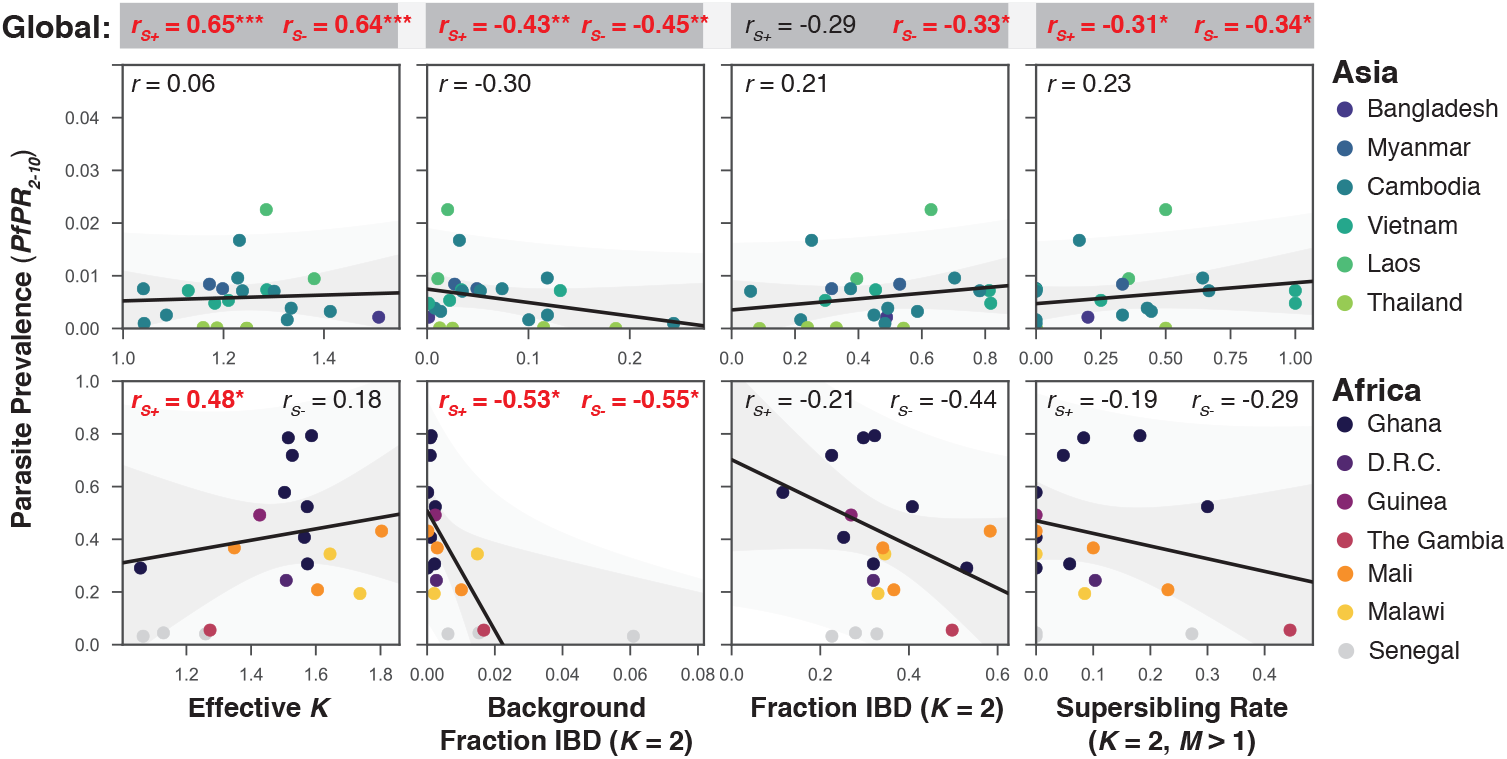
The relationship between *P. falciparum* prevalence and characteristics of mixed infection. Four mixed infection statistics are shown including the average effective number of strains (Effective k, first column), given by 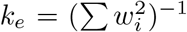, where *w*_*i*_ is the proportion of the *i*th strain; background IBD observed between clonal samples (Background Fraction IBD, second column); fraction IBD within *K* = 2 mixed infections (Fraction IBD, third column); and the rate of *K* = 2 mixed infections classified as having *M >* 1 (Supersibling Rate, fourth column). Each point relates to a row in Table 1 from different sampling locations and years. Pearson’s *r* is computed globally (shown at top in a grey box for each statistic), across Asian countries (upper panel) and across African countries (lower panel). Globally and for Africa, the correlations were computed including Senegal (*r*_*S*+_) and excluding Senegal (*r*_*S*−_). The slope and confidence intervals for the regression line excluding Senegal are drawn. Significant correlations (*p* < 0.05) are highlighted in red and significance levels indicated by asterisks (* < 0.05, ** < 0.01, *** < 0.001)

## 4 Discussion

It has long been appreciated that mixed infections are an integral part of malaria biology. However, determining the number, proportions, and haplotypes of the strains that comprise them has proven a formidable challenge. Previously we developed an algorithm, `DEploid`, for deconvoluting mixed infections (Zhu et al., 2018). However, we subsequently noticed the presence of mixed infections with highly related strains in which the algorithm performed poorly, particularly with low-frequency minor strains. Mixed infections containing highly related strains represent an epidemiological scenario of particular interest, because they are likely to have been produced from a single mosquito bite, itself multiply infected, and in which meiosis has occurred to generate sibling strains. Thus, we developed an enhanced method, `DEploidIBD`, capable not only of deconvoluting highly related mixed infections, but also inferring IBD segments between all pairs of strains present in the infection. We note that limitations and technical difficulties remain, including deconvoluting infections with more than three strains, analysing data with multiple infecting species, coping with low-coverage data, and selecting appropriate reference panels from the growing reference resources.

The application of `DEploidIBD` to the 2,344 samples in the Pf3k project has revealed the extent and structure of relatedness among malaria infections and how these characteristics vary between geographic locations. We found that 1,026 (44%) of all samples in Pf3k were mixed, being comprised of 480 *K* = 2 infections, 372 *K* = 3 and 127 *K* = 4 infections. Across the entire data set, the total number of genomes extracted from mixed infections is nearly double the number extracted from clonal infections (2,584 genomes from *K >* 1 vs. 1,365 from *K* = 1). We also found considerable variation, between countries and continents in the characteristics of mixed infections, suggesting that they are sensitive to local epidemiology. Previous work has highlighted the utility of mixed infection rate in discerning changes in regional prevalence, and we re-enforce that finding here, observing a significant correlation between the effective number of strains and parasite prevalence across Pf3k collection sites. Similarly, using `DEploidIBD` we also observe significant geographical variation in the relatedness profiles of strains within mixed infections. Interestingly, this variation is structured such that regions with high rates of mixed infection tend to contain strains that are less related, resulting in a significant negative correlation between mixed infection rate and mean relatedness within those infections.

The ability to identify the extent and genomic structure of IBD enables inference of the mechanisms by which mixed infections can arise. A mixed infection of *K* strains can be produced by either *K* independent infectious bites or by *j < K* infectious bites. In the first case, parasites are delivered by separate vectors and no meiosis occurs between the distinct strains, thus any IBD observed in the mixed infection must have pre-existed as background IBD between the individual strains. In the second case, meiosis may occur between strains, resulting in long tracts of IBD. The exact amount of IBD produced by meiosis is a random variable, dependent on outcomes of meiotic processes, such as the number of recombination events, the distance between them, and the segregation of chromosomes. Importantly, the mean IBD produced during meiosis in *P. falciparum* also depends on the number and type (selfed vs outbred) of oocysts in the infectious mosquito. Here, we have shown, from first principles, that the amount of IBD expected in a single-bite mixed infection produced from two unrelated parasites strains will always be slightly less than 1*/*2, and potentially as low as 1*/*3 (see Supplementary Materials).

To quantify the distribution of IBD statistics expected through different mechanisms of mixed infection, we developed a Monte Carlo simulation tool, `pf-meiosis`, which we used to infer the recent transmission history of individuals with dual (*K* = 2) infections. We considered mixed infection chains, in which *M* successive rounds of meiosis, transmission to host, and uptake by vector can result in sibling strain infections with very high levels of IBD. Using this approach, we found that 47% of dual infections within the Pf3k Project likely arose through co-transmission events. Moreover, and particularly within Asian population samples, we found evidence for long mixed infection chains (*M >* 1), representing repeated co-transmission without intervening superinfection. This observation is not a product of lower genetic diversity in Asia, as differences in background IBD between countries have been controlled for in the simulations. Rather, it reflects true differences in transmission epidemiology between continents. These findings have three important consequences. Firstly, they suggest that successful establishment of multiple strains through a single infection event is major source of mixed infection. Second, they imply that the bottlenecks imposed at transmission (to host and vector) are relatively weak. Finally, they indicate that the differing mechanisms causing mixed infections reflect aspects of local epidemiology.

We note that a non-trivial fraction (17%) of all mixed infections had patterns of IBD inconsistent with the simulations (typically with slightly higher IBD levels than background but lower than among siblings). We suggest three possible explanations. A first is that the unclassified samples result from the IBD profiles produced by `DEploidIBD`, in particular the overestimation of short IBD tracks, similar to the issue observed by (Wong et al., 2018). Alternatively, our estimate of background IBD, generated by combining pairs of random clonal samples from a given country into an artificial *M* = 0 mixed infection, will underestimate true background IBD if there is very strong local population structure. Finally, we only simulated simple mixed infection transmission chains, at the exclusion of more complex transmission histories, such as those involving strains related at the level of cousins. The extent to which such complex histories can be inferred with certainty remains to be explored.

Lastly, our results show that the rate and relatedness structure of mixed infections correlate with estimated levels of parasite prevalence, at least within Africa, where prevalence is typically high (Smith et al., 1993). In Asia, which has much lower overall prevalence, as well as greater temporal (and possibly spatial) fluctuations, we do not observe such correlations. However, it may well be that other genomic features that we do not consider in this work could provide much higher resolution, in space and time, for capturing changes in prevalence than traditional methods. Testing this hypothesis will lead to a much greater understanding of how genomic data can potentially be used to inform global efforts to control and eradicate malaria.

## 5 Methods and Materials

The data analysed within this paper were collected and made openly available to researchers by member of the Pf3k Consortium. Information about studies within the data set can be found at https://www. malariagen.net/projects/pf3k#sampling-locations. Detailed information about data processing can be found at https://www.malariagen.net/data/pf3k-5. Briefly, field isolates were sequenced to an average read depth of 86 (range 12.6 – 192.5). After removing human-derived reads and mapping to the 3D7 reference genome, variants were called using GATK best practice and approximately one million variant sites were genotyped in each isolate. After filtering samples for low coverage and cross-species contamination, 2,344 samples remained. The Supplementary Material provides details on the filters used and data availability. For deconvolution, samples were grouped into geographical regions by genetic similarity; four in Africa, and three in Asia. (Table 1). Reference panels were constructed from the clonal samples found at each region. Since previous research has uncovered severe population structure in Cambodia (Miotto et al., 2013), we stratified samples into West and North Cambodia when performing analysis at the country level.

## 6 Acknowledgements

This study was supported by the Wellcome Trust (206194, 090770, 204911, 100956/Z/13/Z to GM), the Medical Research Council (G0600718), the UK Department for International Development (M006212) and the Li Ka Shing Foundation (to GM). This study used data from the MalariaGEN Pf3k Project. Genome sequencing was done by the Wellcome Sanger Institute (WSI), and sample collections were coordinated by the MalariaGEN Resource Centre. The samples from Senegal were was supported by funding from the Bill and Melinda Gates Foundation to Dyann Wirth, and sequenced by the Broad Institute. We thank the staff of the WSI Sample Logistics, Sequencing, and Informatics facilities for their contribution; all patients and collaborators contributing samples and data to the Pf3k project. We acknowledge and appreciate the critical role of administrator team and research computing support team, especially Victoria Cornelius for keeping non-scientific business running smoothly.

## 7 Data availability

Metadata on samples is available from ftp://ngs.sanger.ac.uk/production/pf3k/release_5/pf3k_release_ 5_metadata_20170804.txt.gz. Sequence data (aligned to *P*lasmodium falciparum strain 3D7 v3.1 reference genome sequences, for details see ftp://ftp.sanger.ac.uk/pub/project/pathogens/gff3/2015-08/ Pfalciparum.genome.fasta.gz) is available from ftp://ngs.sanger.ac.uk/production/pf3k/release_ 5/5.1/. Diagnostic plots for the deconvolution of all samples can be found at https://github.com/mcveanlab/mixedIBD-Supplement and deconvoluted haplotypes can be accessed at ftp://ngs.sanger.ac. uk/production/pf3k/technical_working/release_5/mixedIBD_paper_haplotypes/. Code implementing the algorithms described in this paper, `DEploidIBD`, is available at https://github.com/DEploid-dev/DEploid. Code to generate *in silico* lab mixture of 4 strains are available at https://github.com/DEploid-dev/ `DEploid`-Data-Benchmark-in_silico_lab_mixed_4s. Code to generate *in silico* field mixtures of 2, 3, 4 strains are available at https://github.com/DEploid-dev/DEploid-Data-Benchmark-in_silico_field.

---

**Figure 1-Figure supplement 1.**
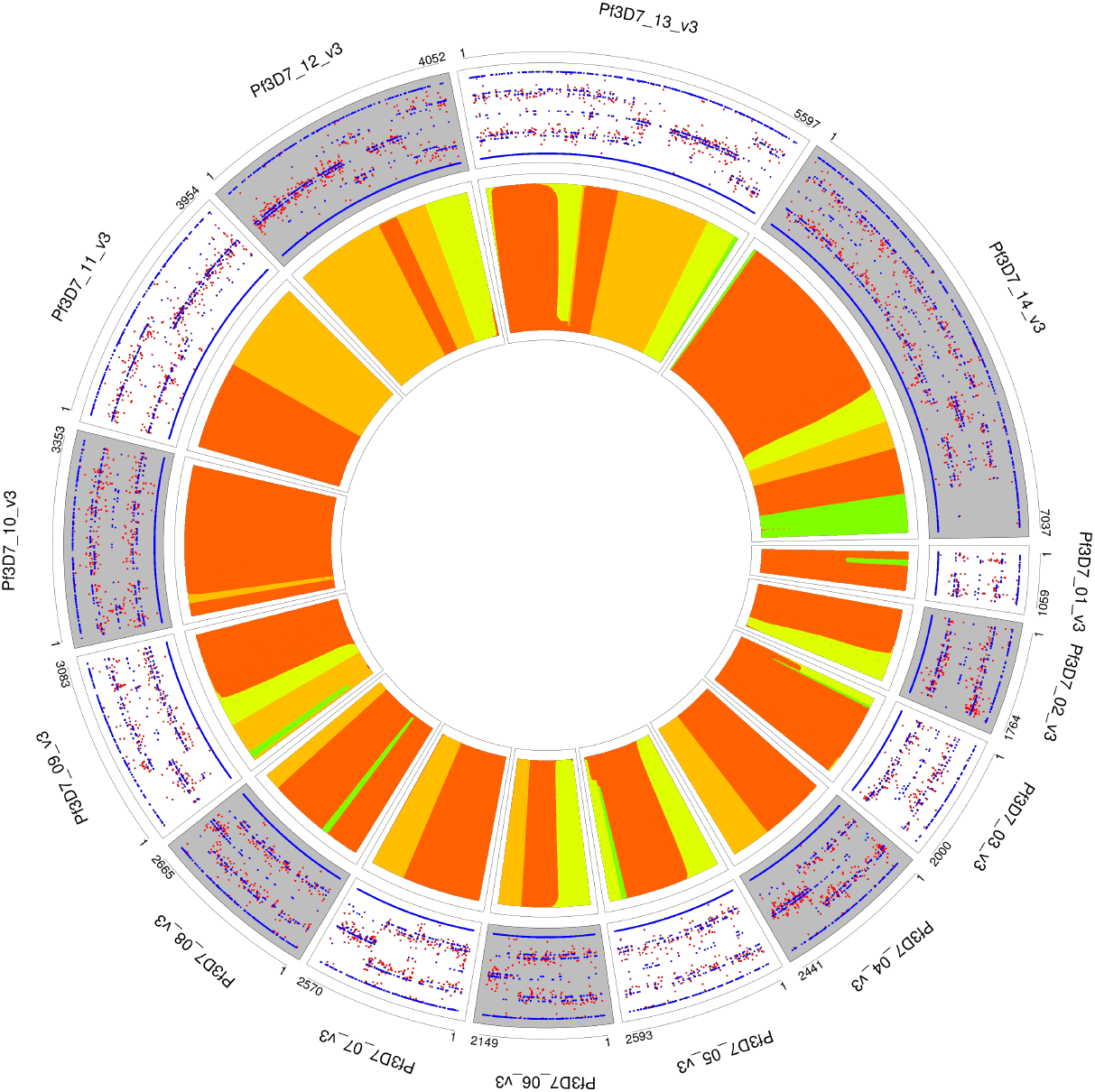
Whole genome deconvolution of field sample PD0577-C. The outer ring shows the expected within-sample allele frequency (WSAF) (blue) and observed WSAF (red) across the genome. Red and blue points indicate observed and expected allele frequencies within the isolate. The inner ring indicates the IBD states among the three strains: green segments indicate where all three strains are IBD; yellow, orange and dark orange segments indicate the regions where one pair of strains are IBD but the others are not. In no region are all three strains inferred to be distinct, suggesting that the three strains are siblings.

**Figure 1-Figure supplement 2.**
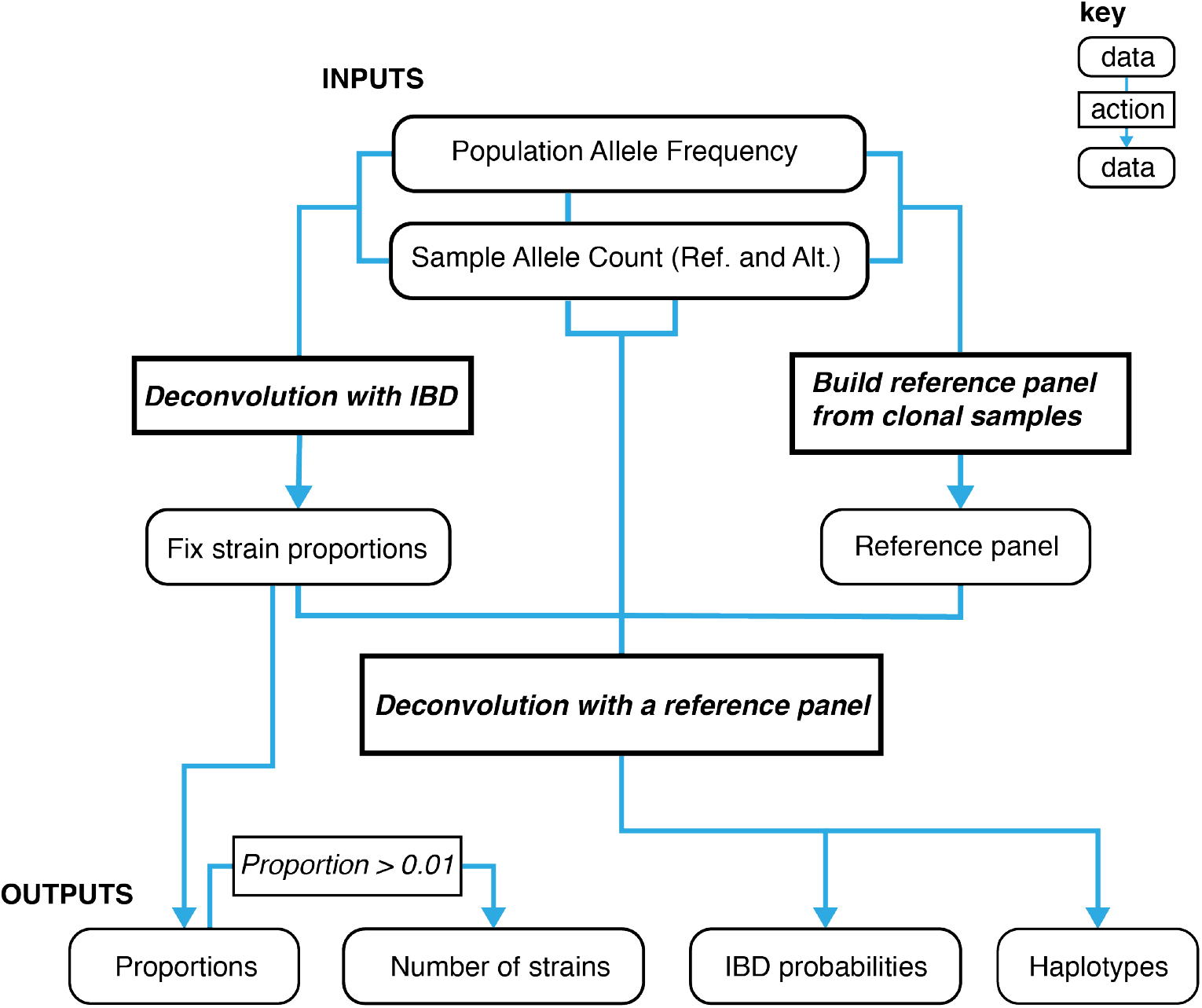
A graphical overview of the data types and work flows for `DEploidIBD`. The boxes at the bottom represent final outputs of the pipeline. The rectangular boxes indicate when `DEploidIBD` is executed, with inputs highlighted by blue arrows. The process has three key steps: Step 1. A reference panel for the set of samples is constructed from high confidence clonal haplotypes, either identified from within a study or from an external resource, such as Pf3k. Step 2: `DEploidIBD`, using population level allele frequencies, is used to infer the number of strains, strain proportions and IBD profile within each sample. Step 3: `DEploidIBD` is re-run on each sample to infer haplotypes, but with the proportions estimated in Step 2 fixed and this time using the haplotype (LD-aware) method previously implemented in `DEploid`.

**Figure 2-Figure supplement 1.**
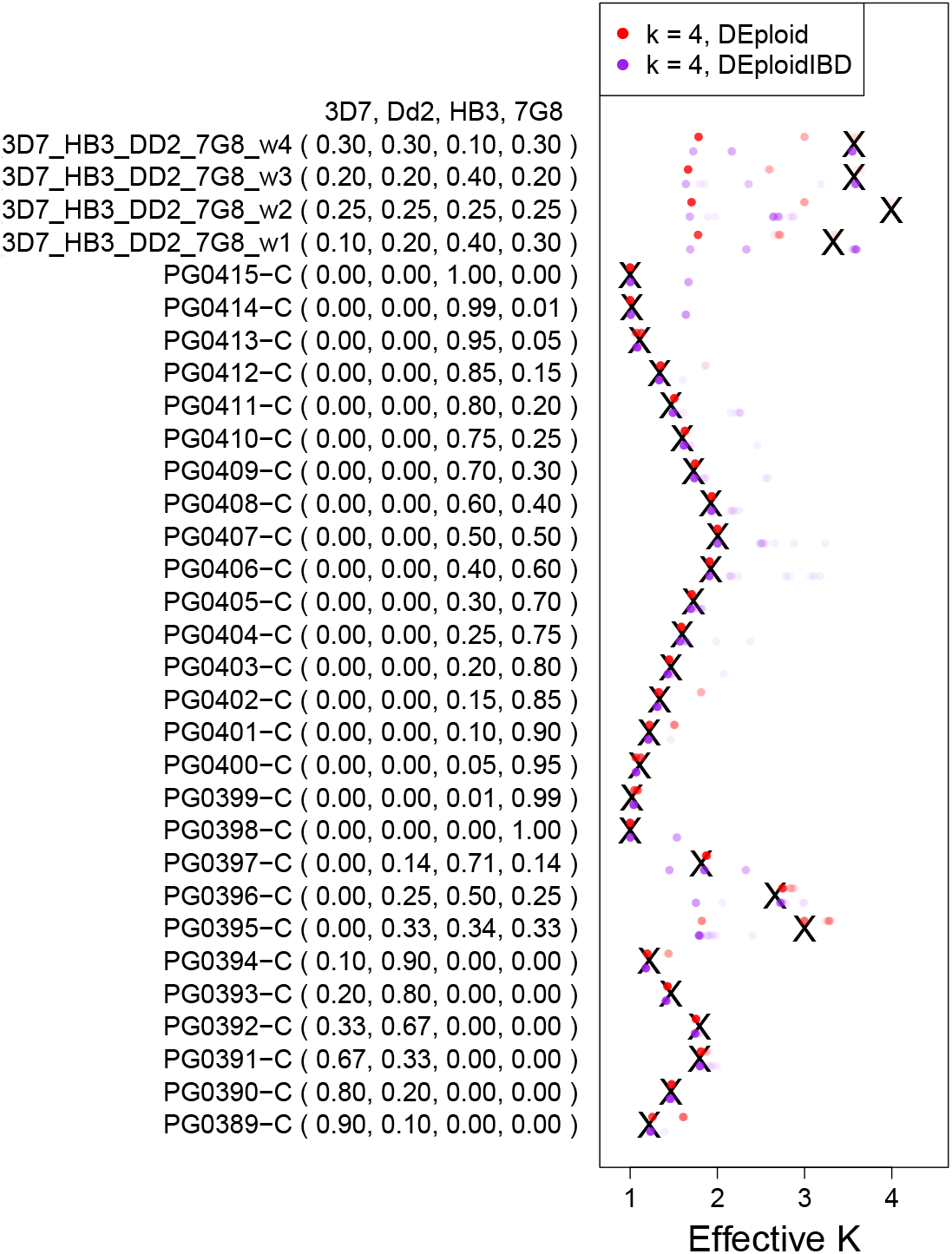
Validation of `DEploidIBD` using 27 *in vitro* lab mixtures and 4 *in silico* mixtures. A reference panel of the laboratory strains (3D7, Dd2, HB3 and 7G8; Panel V) was used to deconvolute samples with `DEploid`. Each experiment is performed with and without IBD inference and with the maximum number of 4 strains. Black crosses indicate the true effective number of strains. Coloured crosses (`DEploid` in red, `DEploidIBD` in purple) indicate median values obtained from 30 replicates using the algorithm indicated in the legend. The coloured dots show the inferred effective number of strains across replicates with intensity proportional to fraction. Note one sample where balanced proportions of three strains results in the LD-free (`DEploid-IBD`) approach fitting the data as a mixture of two strains with proportions of 1/3 and 2/3. For *in silico* mixtures of four strains, `DEploid` performs poorly. `DEploidIBD` shows some improvement in unbalanced mixtures, though misclassifies *K* = 4 mixtures as only having thress strains.

**Figure 2-Figure supplement 2.**
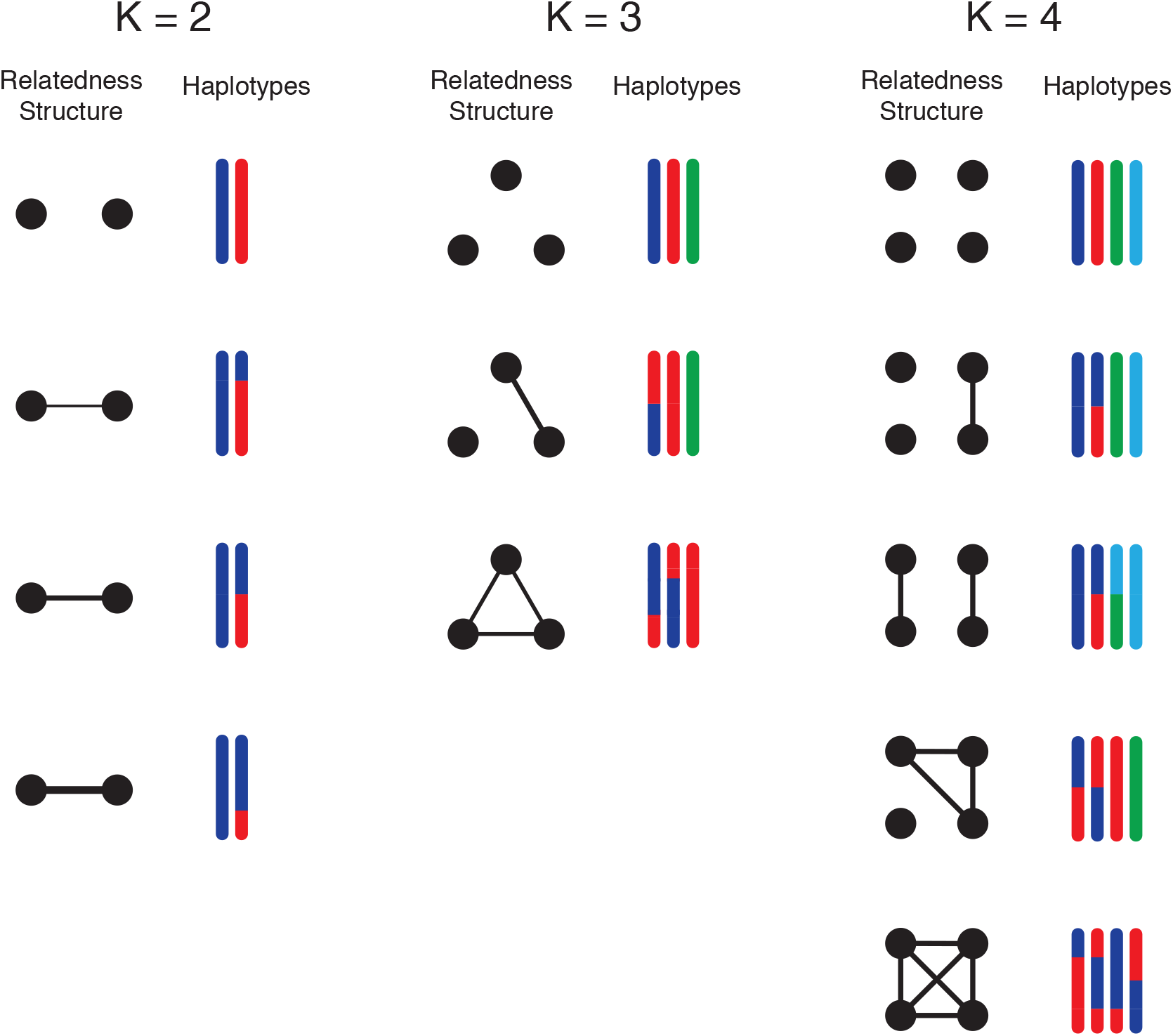
Illustration of simulation study design. We conduct simulation studies to mimic *K*-mixtures (top row) as results of *b*-biting events, where *K* ∈ {2, 3, 4} and 1 ≤ *b* ≤ *K*. For each *K*-mixture, the left column illustrate the overall relationship between strains (black dots): connected dots imply strains are from the same mosquito bite. The level of relatedness between parasite strains is reflected by the haplotype segment copied from the parental strains within the mosquito. Each colour represents a unique strain within the mosquito, which we randomly draw from field clonal haplotypes. For example, when *K* = 2, we consider the case that the two strains are from two independent mosquito bites; on the other hand, when two strains are from the same mosquito bite, we consider scenarios of low (25%), moderate (50%) and high (75%) relatedness between two sibling strains. These events are represented in the second, third and forth rows respectively. For *K* = 3, we consider mixed-infection events as products of three mosquito bites, two mosquito bites and a single bite. For *K* = 4, we consider mixed infections as products of four mosquito bites, three mosquito bites, two mosquito bites and a single bite. We further divide the possibilities of the 2-bite event into the case that both bites pass on two strains (2+2) and the other possibility that one bite passes on a single strain and the other bit passes on three strains (1+3).

**Figure 2-Figure supplement 3.**
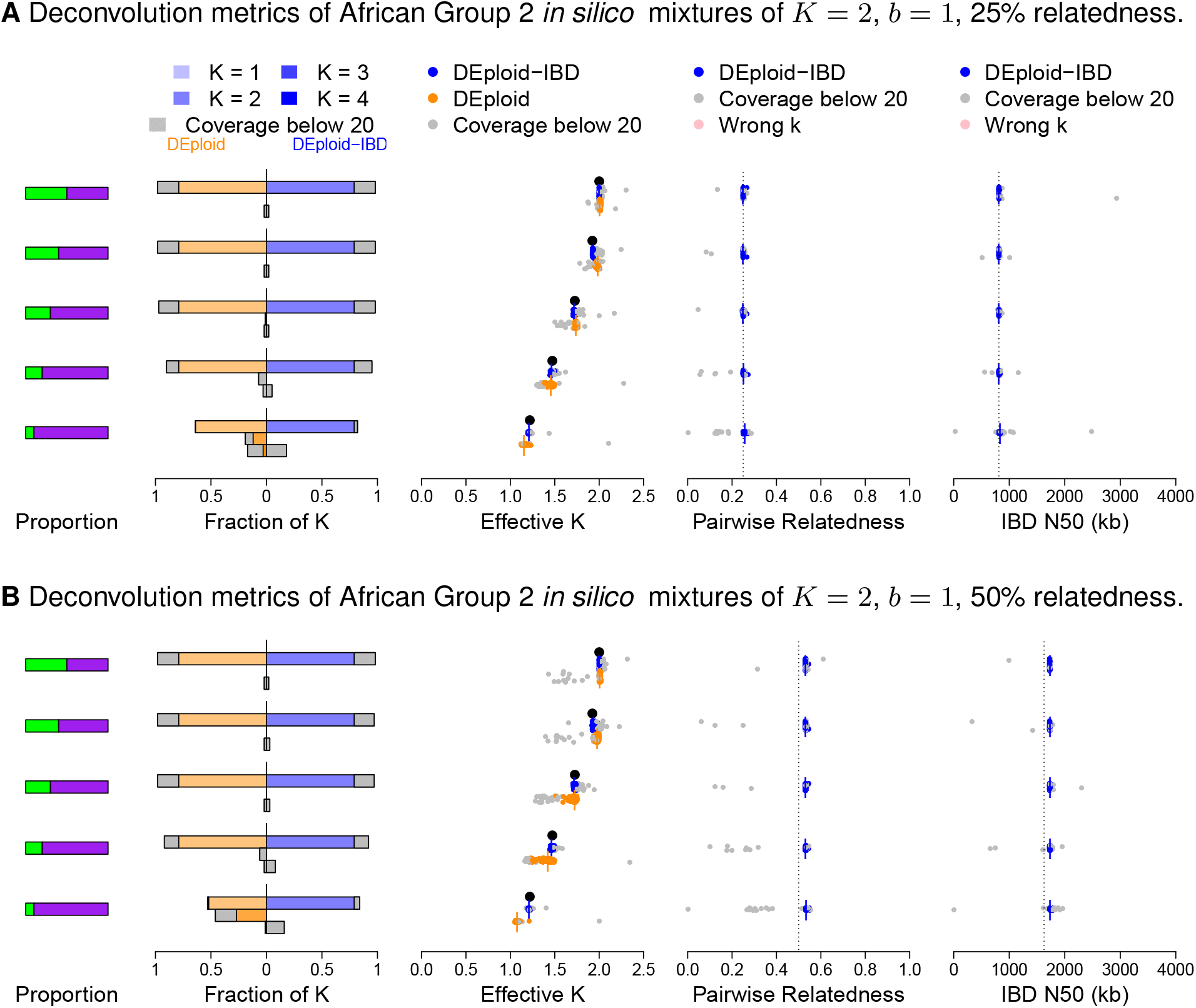
Additional comparison of `DEploidIBD` and `DEploid` on 100 *in silico* mixtures of two strains from Africa with low and moderate relatedness, illustrated by sub panels (A) and (B), respectively. Detailed panel description can be found in the caption to Figure **??**. `DEploid` generally performs well for samples of low within sample relatedness, though struggles when the minor strain proportion is below 30%. In contrast, `DEploidIBD` consitently performs well.

**Figure 2-Figure supplement 4.**
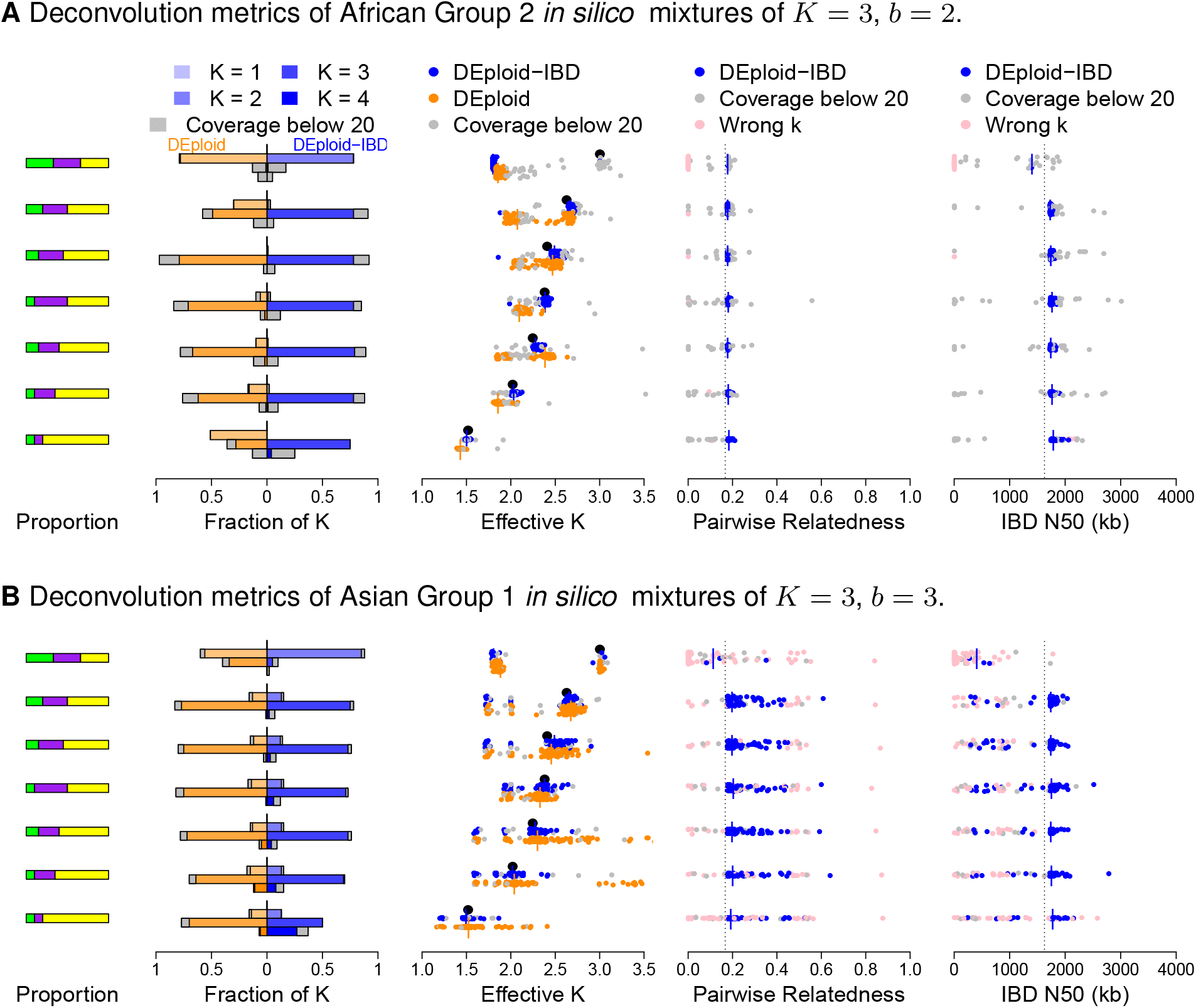
Additional comparison of `DEploidIBD` and `DEploid` on *in silico b* = 2 bite mixtures of *K* = 3 strains from Africa and Asia, illustrated by sub panels (A) and (B), respectively. Detailed panel descriptions can be found in the caption to Figure **??**. The unrelated strain provides a strong signal in allele frequency imbalance for `DEploid` to detect and therefore performs better than dealing with *b* = 1 mixtures. Comparing (A) and (B), pairwise relatedness estimates are noisy in Asia because of the background IBD. However, background relatedness generates shorts segments of IBD and therefore leads to IBD N50 underestimation.

**Figure 2-Figure supplement 5.**
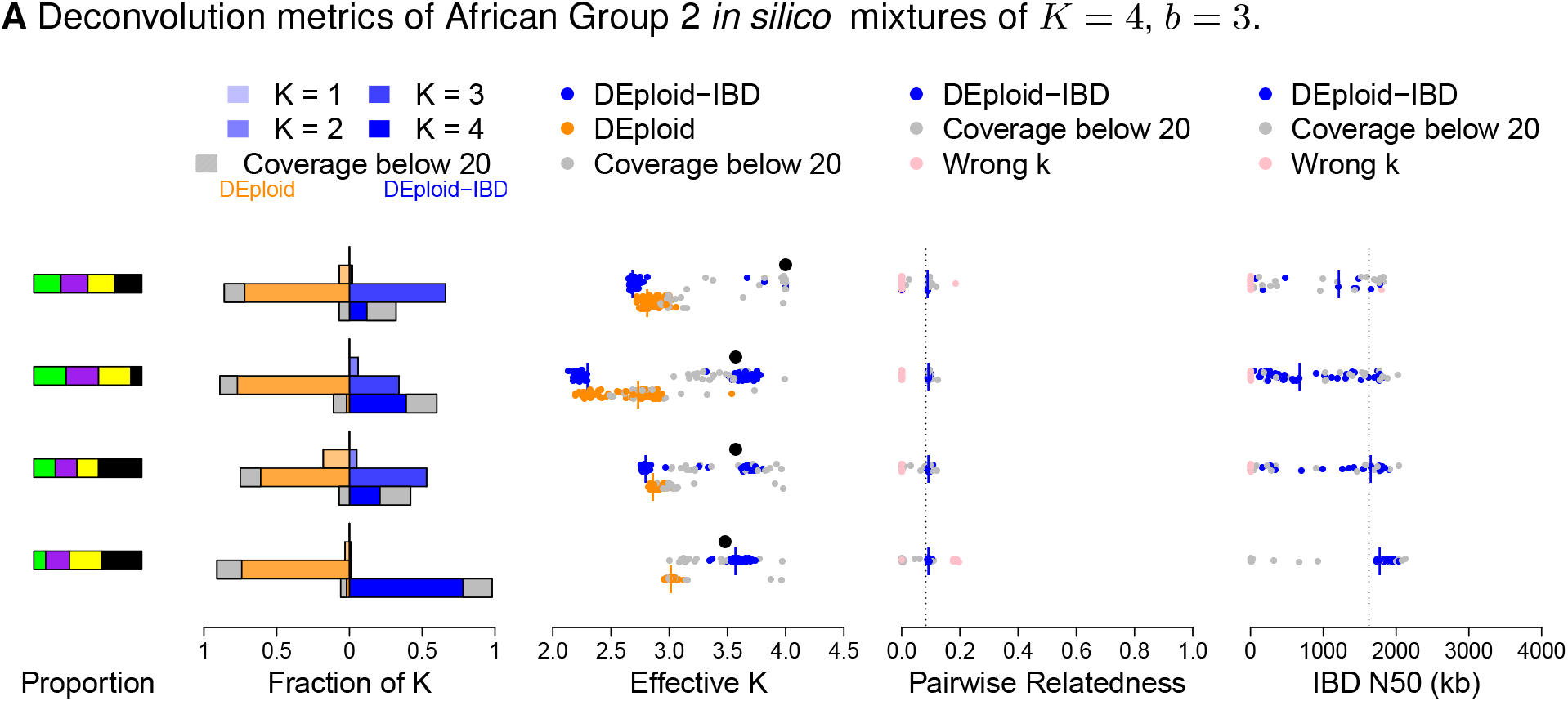
Comparison of `DEploidIBD` and `DEploid` on 100 *in silico b* = 3 bite mixtures of four strains from Africa. Detailed panel descriptions can be found in the caption to Figure **??**. `DEploid` performs poorly in all cases. In contrast, `DEploidIBD` performs well when all four strains have unequal proportions, but is less accurate when some strains have equal proportion.

