## Supplementary material for "The origins and relatedness structure of mixed infections vary with local prevalence of *P. falciparum* malaria"

#### Contents

**S1 Deconvolution in the presence of IBD** **S1-2**
S1.1 Notation . . . S1-2
S1.2 Modelling mixed infections with IBD . . . S1-2
S1.3 Parameter estimation using Markov chain Monte Carlo . . . S1-4
S1.3.1 Metropolis-Hastings update for proportions . . . S1-4
S1.3.2 Gibbs sampling update for haplotype and IBD-configuration update . . . S1-4
S1.4 Gibbs sampling update for IBD parameter . . . S1-5
S1.5 Implementation details . . . S1-5
S1.6 Commands . . . S1-6
S1.7 Error analysis . . . S1-7

**S2 Generating *in silico* mixtures for deconvolution benchmark** **S1-8**
S2.1 *In silico* mixtures of lab strains . . . S1-8
S2.2 *In silico* mixtures of two field strains . . . S1-8
S2.3 *In silico* mixtures of three field strains . . . S1-8
S2.4 *In silico* mixtures of four field strains . . . S1-9

**S3 Pf3k field sample analysis** **S1-10**
S3.1 Sample choice . . . S1-10
S3.2 Data filtering . . . S1-10
S3.2.1 High leverage data points . . . S1-10
S3.3 Analysis preparation . . . S1-11
S3.4 Haplotype quality assessment . . . S1-12
S3.5 Combining clonal sample pairs for background IBD computation . . . S1-13

**S4 Expected levels of IBD in *P.falciparum* mixed infections** **S1-15**

### S1 Deconvolution in the presence of IBD

#### S1.1 Notation

We use the same notations as Zhu et al. (2018) (see Table S1.1). Our data, $D$, are the allele read counts of sample $j$ at a given site $i$, denoted as $r_{j,i}$ and $a_{j,i}$ for reference (REF) and alternative (ALT) alleles respectively. These have assigned values of 0 and 1, respectively. Here we consider only bi-allelic loci, though future extensions to the model to include multi-allelic sites could be accommodated. The empirical allele frequencies within a sample (WSAF) $p_{j,i}$ and at population level (PLAF) $f_i$ are calculated by $\frac{a_{j,i}}{a_{j,i}+r_{j,i}}$ and $\frac{\sum_j a_{j,i}}{\sum_j a_{j,i}+\sum_j r_{j,i}}$ respectively. Since all data in this section refers to the same sample, we drop the subscript $j$ from now on.

| | |
|---|---|
| $i$ | Marker index |
| $j$ | Sample index |
| $r$ | Read count for reference allele |
| $a$ | Read count for alternative allele |
| $f$ | Population level allele frequency (PLAF) |
| $K$ | Number of distinct strains within sample |
| $l$ | Number of sites |
| $\mathbf{w}$ | Proportions of strains |
| $\mathbf{x}$ | Log titre of strains |
| $\mathbf{h}_i$ | Allelic states of $K$ parasite strains at site $i$ |
| $h_{k,i}$ | Allelic state of parasite strain $k$ at site $i$ |
| $p$ | Observed within sample allele frequency (WSAF) |
| $q$ | Unadjusted expected WSAF |
| $\pi$ | Adjusted expected WSAF |
| $e$ | Probability of read error |
| $\mathcal{S}_i$ | IBD configuration at site $i$ |
| $\theta$ | Probability of non-IBD in a mixture of two strains |

Table S1.1: Notation used in this article.

#### S1.2 Modelling mixed infections with IBD

We describe the mixed infection problem by considering the number of strains, $K$, the relative abundance of each strain, $\mathbf{w}$, and the allelic state of the $k$th strain at the $i$th locus, $\mathbf{h}_{k,i}$. In addition to Zhu et al. (2018), we also infer the identity-by-descent (IBD) state $\mathcal{S}_i$, which describes the strain relationship with each other at site $i$. For example, for three strains, the IBD-state could be one of the five cases: (1) all strains are not IBD; (2) strains 1 and 2 are IBD; (3) strains 1 and 3 are IBD; (4) strains 2 and 3 are IBD; (5) all strains are IBD (see Table S1.2). To simplify the problem, we assume independence between markers, and drop the subscript $i$ from now on. As previously, we use a Bayesian approach and define the posterior probabilities of $K$, $\mathbf{w}$, $\mathbf{h}$ and $\mathcal{S}$, as

$$P(K,\mathbf{w},\mathbf{h},\mathcal{S}|e,D) \propto L(K,\mathbf{w},\mathbf{h},\mathcal{S}|e,D) \times P(K,\mathbf{w},\mathbf{h},\mathcal{S}), \tag{S1.1}$$

where $e$ is the read error rate. Moreover, we can decompose the joint prior as

$$P(K,\mathbf{w},\mathbf{h},\mathcal{S}) = P(K) \times P(\mathbf{w}|K) \times P(\mathcal{S}|K) \times P(\mathbf{h}|\mathcal{S}). \tag{S1.2}$$

The priors on the number of strains, $K$, and their proportions, $\mathbf{w}$, are as in Zhu et al. (2018), with associated hyperparameters being user-defined. To model IBD configurations and resulting haplotypes we introduce a

| Index | IBD state | | |
|---|---|---|---|
| | K = 2 | K = 3 | K = 4 |
| 0 | 0-1 | 0-1-2 | 0-1-2-3 |
| 1 | 0-0 | 0-0-1 | 0-0-1-2 |
| 2 | | 0-1-0 | 0-1-0-2 |
| 3 | | 0-1-1 | 0-1-2-0 |
| 4 | | 0-0-0 | 0-1-1-2 |
| 5 | | | 0-1-2-1 |
| 6 | | | 0-1-2-2 |
| 7 | | | 0-0-1-1 |
| 8 | | | 0-1-0-1 |
| 9 | | | 0-1-1-0 |
| 10 | | | 0-0-0-1 |
| 11 | | | 0-0-1-0 |
| 12 | | | 0-1-0-0 |
| 13 | | | 0-1-1-1 |
| 14 | | | 0-0-0-0 |

Table S1.2: IBD configurations for two, three and four strains, ordered top to bottom by the number of IBD pairs. The (zero-indexed) notation indicates the type assigned to each haplotype, thus 0-1 indicates non-IBD for two strains, while 0-1-2-2 indicates four strains in which the third and fourth are IBD.

parameter $\theta$, which is the probability of outbreeding (i.e. non-IBD) for a mixed infection with two strains. Specifically, for $K$ strains, the prior probability on configuration $\mathcal{S}$ is a function of the number of distinct strains in the configuration, $C_{\mathcal{S}}$, and the number of distinct configurations with the same number of distinct strains, $M_{\mathcal{C}}$:

$$P(\mathcal{S}|K) = \binom{K-1}{C_{\mathcal{S}}-1} \frac{\theta^{C_{\mathcal{S}}-1}(1-\theta)^{K-C_{\mathcal{S}}+1}}{M_{\mathcal{C}}}. \tag{S1.3}$$

That is, the number of distinct strains is drawn from a binomial distribution with parameters $K$ and $\theta$ and the IBD configuration is uniformly drawn from among those with the same number of distinct strains. Consequently, the expected number of distinct haplotypes (i.e. non-IBD) in an infection with $K$ strains is $1+\theta(K-1)$. The value of $\theta$ estimated within the Markov chain Monte Carlo (see details below).

To model the transitions between IBD states we make the approximation that the IBD state at the $i+1$th site is the same as at the $i$th site, with probability $1-p_{rec}$ and, if there is a recombination event, the IBD state is drawn from the stationary distribution described above.

To model allelic states (or genotype), conditional on IBD state, we assume that allelic states for each IBD group are independent Bernoulli draws given the population allele frequency (PLAF). For example, if $K = 3$ and only strains 1 and 2 are IBD (state 0,0,1), the genotype at site $i$, $\mathbf{h_i}$, could be $\{0,0,0\}$; $\{0,0,1\}$; $\{1,1,0\}$ or $\{1,1,1\}$, with probabilities $(1-f)^2$, $(1-f)f$, $f(1-f)$ and $f^2$ respectively. The joint prior on IBD and allelic states is referred to as $\psi(\mathcal{S},\mathbf{h})$. Note that we ignore association (known as linkage disequilibrium or LD) between nearby alleles, though note that in implementation we run `DEploidIBD` twice; first as described here to estimate strain number and proportions, allowing for IBD, then subsequently with a reference panel, as for `DEploid`, but with the strain number and proportions fixed. This latter step does model LD and consequently results in better estimates of strain haplotypes.

Details of the emission model are the same as described in Zhu et al. (2018). Briefly, the reference and alternative allele read counts at each site are modelled as being drawn from a beta-binomial distribution given the expected WSAF, $q = \mathbf{w}^T\mathbf{h}$. To incorporate sequencing error, we modify the expected WSAF such that allele are misread with probability $e$. Thus, the adjusted expected WSAF becomes

$$\pi_i = q_i + (1-q_i)e - q_i e = q_i + (1-2q_i)e. \tag{S1.4}$$

As previously, we model over-dispersion in read counts relative to the Binomial using a Beta-binomial distribution.

#### S1.3 Parameter estimation using Markov chain Monte Carlo

To infer the relatedness between strains within in a mixed infection and their relative proportions we use a Markov chain Monte Carlo (MCMC) approach. We learn the relative abundance of each strain by exploiting signatures of within-sample allele frequency imbalance, using a Metropolis-Hastings algorithm, which samples proportions, $\mathbf{w}$, given IBD-configurations, $\mathcal{S}$. We then use a Gibbs sampler to update $\mathcal{S}$ and $\mathbf{h}$ for a given $\mathbf{w}$ by first sampling the IBD state from the posterior, and then sampling the allelic configuration (genotype) from the selected IBD state at each site to update haplotypes. Note that the hidden Markov model structure for IBD states along the chromosome leads to efficient algorithms for calculating quantities of importance. Notably, by summing over allelic configurations that are consistent with a given IBD configuration, we can - for a given set of strain proportions - calculate the likelihood integrated over all possible IBD configurations, which leads to efficient sampling.

##### S1.3.1 Metropolis-Hastings update for proportions

We update strain proportions, $\mathbf{w}$, through modelling underlying log titres, $\mathbf{x}$, where $w_i \propto exp(x_i)$. As previously, log titres are assumed to be drawn from a $N(-log(K), \sigma^2)$ distribution under the prior. To update, we choose $i$ uniformly from $K$ and propose new $x_i'$s from $x_i' = x_i + \delta x$, where $\delta x \sim N(0, \sigma^2/s)$, and $s$ is a scaling factor. The new proposed proportion is therefore $\frac{\exp(x_i')}{\sum_{k=1}^{K} \exp(x_k')}$. Since the proposal distribution is symmetrical, the Hastings ratio is 1. The proposed update is accepted with probability

$$\min\left(1, \frac{P(\mathbf{w}'|K)}{P(\mathbf{w}|K)} \frac{L(\mathbf{w}', \mathbf{h}|e, D)}{L(\mathbf{w}, \mathbf{h}, |e, D)}\right).$$

Note that, conditional on the current estimate of strain haplotypes, $\mathbf{h}$, the likelihood is not a function of the specific IBD configuration.

##### S1.3.2 Gibbs sampling update for haplotype and IBD-configuration update

As described above, in `DEploidIBD`, allelic states at different sites within a haplotype are independent. However, IBD states are connected through a Markov process. We therefore update the IBD configuration and strain haplotypes in a two stage process. First, we use the Forward algorithm to first compute the integrated likelihood - that is the probability of observing the read-level data integrating over all possible IBD configurations and allelic states. Defining $F_{i,\mathcal{S},\mathbf{h}}$ to be the integrated likelihood for the set of paths ending in IBD configuration $\mathcal{S}$ at site $i$ and with allelic configuration $\mathbf{h}$, it follows that

$$F_{i,\mathcal{S},\mathbf{h}} = P(D_i|e, \mathbf{h})\big[(1 - p_{rec})F_{i-1,\mathcal{S},\mathbf{h}} + \psi(\mathcal{S}, \mathbf{h})p_{rec} \sum_{m,n} F_{i-1,m,n}\big], \tag{S1.5}$$

where $\psi(\mathcal{S}, \mathbf{h})$ is the prior on the combination of IBD configuration $\mathcal{S}$ and allelic configuration $\mathbf{h}$ as defined in Section S1.2. Note that the summation term is identical for all IBD / allelic configurations.

By storing the output of the Forward algorithm we can then sample from the posterior distribution of the IBD- and allelic-configurations (given current proportions). That is, we sample $\mathcal{S}_l, \mathbf{h}_l \propto F_{l,\mathcal{S},\mathbf{h}}$, and then previous steps proportion to $F_{i,\mathcal{S},\mathbf{h}}$ times the probability to transition to the sampled state at position $i + 1$.

**Caveat: identifiability with balanced mixing**

Using a reference panel free approach means that it can be difficult or impossible to deconvolute samples containing strains with equal proportions. For example, it is impossible to distinguish between $\{\frac{1}{3}, \frac{1}{3}, \frac{1}{3}\}$ and $\{\frac{1}{3}, \frac{2}{3}\}$, or $\{\frac{1}{4}, \frac{1}{4}, \frac{1}{4}, \frac{1}{4}\}$ and $\{\frac{1}{4}, \frac{1}{4}, \frac{1}{2}\}$. Because of the prior we use, the model will prefer to merge haplotypes when possible, which can lead to underestimation of the number of strains. We advise users to apply `DEploid` with multiple runs with and without the '-ibd' flag and see if such problem occurs.

#### S1.4 Gibbs sampling update for IBD parameter

Given the sampled IBD configuration, the IBD parameter $\theta$ can be updated using Gibbs sampling. We first identify the path of the $D$ distinct IBD configurations along the chromosome and, for each, identify the number of distinct IBD groups, $\mathcal{C}_d$. For example, if the IBD state is 0,1,2,2 there are three distinct IBD groups (0,1,2; $\mathcal{C}_d = 3$), while if the IBD state is 0,0,1,1 there are two IBD groups (0,1; $\mathcal{C}_d = 2$). Under the prior, the number of distinct strains (minus 1) is drawn from a binomial distribution with parameters $K-1$ and $\theta$. With a uniform prior on $\theta$, a new value, $\theta'$ is therefore drawn from:

$$\theta' \sim Beta\big(\mathcal{C}_D - D + 1, DK - \mathcal{C}_D + 1\big),$$

where $\mathcal{C}_D = \sum_{d=1}^{D} \mathcal{C}_d$.

#### S1.5 Implementation details

Below we detail a number of implementation details.

- **Approximating the likelihood surface** At any site, the likelihood for the WSAF, $q$, induced by the allelic configuration and strain proportions derives from the beta-binomial model as implemented in `DEploid`. That is, the likelihood of $q_i$ is

$$L(q_i|e, D) \propto \frac{B(a_i + \pi c, r_i + (1-\pi)c)}{B(\pi c, (1-\pi)c)}, \tag{S1.6}$$

  where $\pi_i$ is the adjusted WSAF $\pi = q(1-e) + (1-q)e$ and $c^{-1}$ determines the magnitude of over-dispersion relative to the binomial, with $c \to \infty$ recovering the binomial. Here, we use $c = 100$.

  For computational efficiency, rather than recomputing for every value of $q$, we first approximation the likelihood function for each site through a Beta distribution, matching the first two moments of the posterior on $q$ implied by Equation S1.6 within a uniform prior on $q$. The estimated parameters of the Beta distribution are then used to approximate the likelihood surface in all subsequent calculations.

- **MCMC parameters for deconvolution**

  – **Number of strains**. As described above, we aim to infer more strains than are actually present, starting the MCMC chain with a fixed $K$ (default of 5). In our experience, we find poor performance for $K > 4$, hence use use the flag `-k 4` to specify the number of strains as 4. At the point of reporting, we keep strains with a proportion above a fixed threshold, typically 0.01.

  – **Parameters**. We typically set the log-titre variance $\sigma^2 = 5$, which can be adjusted when working with extremely unbalanced samples (see Zhu et al. (2018) Supplementary Material). We set the read error rate as 0.01 and the rate of mis-copying as 0.01.

  – **Recombination rate and scaling**. We assume a uniform recombination map, where the genetic distance between loci $i$ and $i+1$ is computed by $\gamma_i = D_i/d_m$ where $D_i$ denotes the physical distance between loci $i$ and $i+1$ in nucleotides and $d_m$ denotes the average recombination rate in Morgans bp$^{-1}$. We use the recombination rate for *P. falciparum* of 13,500 base pairs per centiMorgan as reported by Miles et al. (2016). The recombination rate is scaled by a factor $G$, which reflects the effective population size, rate of inbreeding, and relatedness of the reference panel. In practice, we deconvolute over 1 million markers in field samples. We tuned the parameter $G$ using *in vitro* lab mixtures, finding that a value of $G = 20$ ensures small recombination probabilities between any two markers, with a mean of 0.015. A large value of $G$ relaxes the reference panel constraint, becoming an LD free model when $G$ is infinity. The scaled genetic distance $G\gamma$ is used to compute the transition probability of switching from copying reference haplotype $a$ to reference haplotype $b$. Given LD varies enormously between *P. falciparum* populations, we will investigate how to tune this parameter for future improvement. In the current release, our program allows users to apply the flag `-G` to specify a specific value. We note that IBD deconvolution from error-prone data can benefit from even smaller values of $G$, such as 0.1.

  - **Reporting**. We aim to provide users with a single point estimate of the haplotypes, their proportions and IBD configurations, although the full chain is also available for analysis. To achieve this we report values at the last iteration - i.e. we report a single sample from the posterior. However, to measure robustness, we typically repeat the deconvolution with multiple random starting points. We use a majority vote rule on the inferred number of strains; we then select the chain with the lowest average deviance (after removing the burn-in) as our estimate. The deviance measures the difference in log likelihood between the fitted and saturated models, the latter being inferred by setting the WSAF to that of the observed values. These parameters can be modified by users to achieve a preferred balance between computational speed and confidence. By default, we set the MCMC sampling rate as 5, with the first 50% of samples removed as burn in and 800 samples used for estimation.
  - **Reference panel construction**. To infer clonal samples for the reference panel we use the Pf3k project data, running the algorithm without LD on all samples and identifying those with a dominant haplotype (proportion $> 0.99$) as clonal. These clonal samples are grouped by region of sampling to form location-specific reference panels. In addition, we have included a number of reference strains, described in more detail below.

- **Summarising pairwise IBD**

  At the end of a run, we obtain the maximum likelihood estimate of IBD configurations along the chromosome using the Viterbi algorithm. Patterns of IBD are then obtained for all pairs of strains and summarised through the mean fraction IBD and the N50 IBD tract, obtained by identifying all blocks of IBD, sorting them in decreasing size and finding the size of the block such that 50% of all sites that are in IBD lie in blocks of at least this size. A larger N50 statistic indicates more recent common ancestry.

  We note that it is also possible to obtain estimates of pairwise IBD from the full posterior distribution of pairwise IBD (using standard hidden Markov model practices). Typically these give very similar answers to the Viterbi solution, though are typically slightly larger due to the identification of low certainty regions. The posterior mean IBD between pairs is also provided to users as output.

#### S1.6 Commands

Each isolate deconvolution is repeated 15 times with a random seeds of choice from 1 to 15. For each run we obtain an estimate of the number of strains and we take the modal value to be the estimate. For reporting, we use the first run that estimated the consensus number of strains. The text below gives an example of an input script for deconvolution. Full details are available in the documentation at the Github page: https://github.com/mcveanlab/DEploid/.

```
ref=PD0577-C_ref.txt
alt=PD0577-C_alt.txt
plaf=asiaGroup1_PLAF.txt
panel=asiaGroup1PanelMostDiverse10.csv
exludeAt=asiaGroup1_and_pf3k_bad_snp_in_at_least_50_samples.txt

prefix=PD0577-C_IBD
time dEploid -ref ${ref} -alt ${alt} -plaf ${plaf} -panel ${panel} -exclude ${exludeAt} -o
    ${prefix} -nSample 250 -rate 8 -burn 0.67 -ibd -k 4
interpretDEploid.r -ref ${ref} -alt ${alt} -plaf ${plaf} -o ${prefix} -dEprefix ${prefix} -exclude
    ${exludeAt}
```

#### S1.7 Error analysis

For haplotype quality analysis, we compared the inferred haplotype (`DEploid` output) with the true haplotype. In addition to mismatches between the testing haplotype and the truth, the deconvolution process also introduces switch errors, where the haplotypes are correct, but have undergone *in silico* recombination events. In addition, when one strain has low frequency, there might be insufficient data to make accurate inference, resulting in missing segments of the true haplotypes. We refer to this as dropout error. Dropout error can also be caused by the sequencing process, when parts of the genome are not well sequenced (low read count).

When comparing inferred haplotypes to true haplotypes we use a dynamic programming approach to find a description of the differences, optimising a cost function in which switch errors are twice as costly as mismatches and allele dropouts. Code for performing the optimal description, `errorAnalysis.r`, is available at the Github page: https://github.com/DEploid-dev/DEploid-Utilities/. An example analysis for three strains is shown in Figure S1.1.

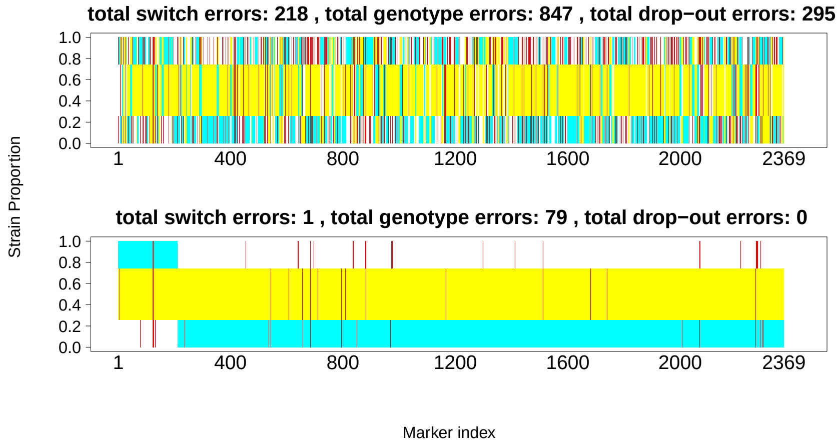


Figure S1.1: Comparison of true and inferred haplotypes for Chromosome 14 (2,369 SNPs) in the lab strain mixture sample PG0396-C after running `DEploidIBD` to infer strain number and proportions (top) and after subsequent refinement of haplotypes by running `DEploid` with Reference Panel V (bottom). The yellow, cyan and white backgrounds identify the haplotype segments from strains 7G8, HB3 and Dd2 respectively. Numbers in the titles indicate the inferred switch, mismatch and dropout errors identified by the dynamic programming approach, with the cost of switch errors being twice that of other errors.

### S2 Generating *in silico* mixtures for deconvolution benchmark

#### S2.1 *In silico* mixtures of lab strains

For *in silico* lab mixtures, we used the lab strain GB4 to provide a whole-genome read depth profile. We drew Bernoulli variables to simulate alternative allele counts, with the probability of success as the adjusted within sample allele frequency at each position (Equation S1.4). The expected WSAF is the dot product of the genotype of the lab strains: 3D7, Dd2, HB3, 7G8 and the proportions. For example, if the genotype is 0, 0, 1, 1 at an arbitrary position, and the mixture proportion of 0.1, 0.2, 0.4 and 0.3 will lead to an expected WSAF of 0.7. We then further adjust the WSAF with an error term (0.01) to account for all technical error/noise: $0.7 \times (1 - 0.01) + (1 - 0.7) \times .01 = 0.696$.

To test the accuracy of `DEploidIBD` in a more realistic setting, we created *in silico* mixtures of 2, 3 and 4 strains given different transmission scenarios from mosquito bites. Let $K$ denote the number strains within each sample, and $b$ denote the number of independent mosquito bites. In such a scenario, a mixed infection containing $K$ strains can be treated as a result of a $b$-bite events, where $K \in \{2, 3, 4\}$ and $1 \leq b \leq K$. For example, a $K = 2$ mixed infection can result from co-transmission, where two strains are passed in a single bite ($b = 1$); or by superinfection, where each strain is delivered by a unique bite ($b = 2$). A mixed infection containing 3 strains is more complex: besides co-infection (3 strains in a single bite, $b = 1$) and superinfection (3 strains in 3 bites, $b = 3$), we can also have a super-infection scenario made up from a co-infection plus a clonal-infection ($b = 2$). The complete simulation procedure (code) is available at https://github.com/DEploid-dev/DEploid-Data-Benchmark-in_silico_field.

#### S2.2 *In silico* mixtures of two field strains

To simulate a mixture of two strains, we randomly selected two strains from 189 clonal samples of African origin (proportions ranging from 10/90% to 50/50%) using Chromosome 14 data. A further 20 randomly chosen samples were used as the reference panel. In order to compare the accuracy of the two methods at different levels of relatedness, we set 0%, 25%, 50% and 75% of the second haplotype the same as the first haplotype to mimic scenarios of unrelated, low, medium and high relatedness respectively. This operation sets a lower limit to the relatedness between two strains, as background relatedness may also exist. We used empirical read depths and drew read counts for the two alleles from binomial proportions (the same approach for generating *in silico* lab mixtures). We excluded sites for analysis at zero alternative allele counts in both targeted samples and reference panel, kept and analysed around 7,757 polymorphic sites (standard deviation 178) for each *in silico* samples.

We repeated the *in silico* experiment with mixtures of two strains from 204 clonal Asian samples, also with mixing proportions of ranging from 10/90% to 50/50%, using about 3,041 sites (s.d. 227) from Chromosome 14.

#### S2.3 *In silico* mixtures of three field strains

We further extended benchmarking to *in silico* mixtures of three strains in African populations with mixing proportions of 10/10/80%, 10/25/65%, 15/25/60%, 10/40/50%, 15/30/55%, 20/30/50%, 33/33/34%. Two generate $b = 1$ infections with three strains, we randomly selected two strains, namely parent A and parent B, from 189 clonal samples of Africa origin, and set the first 33% of the first haplotype and the last 66% of the second haplotype the same as parent A; the rest of the first and second haplotypes and the third haplotype the same as parent B, to mimic the scenario of a 1/3 pairwise relatedness within sample.

In addition to the basic co-infection and super-infection scenarios of 3 strains, we also consider more complex events when co-infection presents within a super-infection. This data simulation is similar to simulating 2 strains of 50% relatedness, with one additional unrelated haplotype. Therefore, the overall pairwise relatedness is 1/2 distributed over three possible pairs, which leads to 1/6.

#### S2.4 *In silico* mixtures of four field strains

We tested *In silico* deconvolution experiments on mixtures of 4 strains. From previous experiments, we have learned that strain compositions with even proportions are difficult to deconvolute. In this set of simulations, we experimented with various unbalanced and balanced proportions including 11/22/30/37%, 25/25/25/25%, 20/20/20/40% and 30/30/30/10%. For some cases, the data generation procedures can easily be modified from previous experiment: a $b = 4$ event of for four strains is equivalent to a 3-bite event of 3 strains with one extra random haplotype; a $b = 3$ event of four strains is equivalent to a 2-bite event of 3 strains with one extra random haplotype. For 2-bite event of 4 strains, there are two possibilities: (i) both bites pass on 2 strains, which is essentially repeating $b = 1$ event of 2 strains twice or (ii) three strains in one bite and one in another, which is essentially a 1-bite event of 3 strains with one extra random haplotype.

### S3 Pf3k field sample analysis

#### S3.1 Sample choice

We used 2,640 samples from the Pf3k project (see https://www.malariagen.net/projects/pf3k). We excluded Nigerian samples from downstream analysis as there were only 5 samples from this country. We discarded samples containing mixed malaria species and those where sequencing coverage depth was below 30X or in which less than 30% of sites were callable. Finally, we exclude all lab strains (including reference strains, artificial mixtures, and crosses samples) and duplicated samples. In total, we retained 2,344 field samples from 13 countries.

#### S3.2 Data filtering

We ran `DEploid-IBD` on high quality biallelic SNPs (both coding and non-coding, tagged with PASS at the QUAL column in the VCF file) from Pf3k (Pf3k Consortium, 2016). Before the additional filtering step described below, this set contained 1,057,830 SNPs.

```
bcftools view \
--include 'FILTER="PASS"' \
--min-alleles 2 \
--max-alleles 2 \
--types snps \
--output-file SNP_INDEL_Pf3D7_01_v3.high_quality_biallelic_snps.vcf.gz \
--output-type z \
SNP_INDEL_Pf3D7_01_v3.combined.filtered.vcf.gz
```

##### S3.2.1 High leverage data points

We found that markers with high coverage for both alleles could mislead our model, inducing it to fit additional strains. We used a threshold of $\geq$ 99.5% coverage (default) to identify markers with extremely high allele counts. We further expanded this list of potential problematic markers by considering their nearest 10 neighbours on both flanks, and excluding those that were tagged more than once (see Figure S3.2). These poorly-genotyped variants are likely to be errors of mapping and genotype calling.

To track down the causes of high leverage points, we assessed nucleotide diversity in the *P. falciparum* genome. We used clonal haplotypes to compute nucleotide diversity by running a sliding window along the genome. At each SNP, we use $n_0$ and $n_1$ to denote counts for reference and alternative alleles, respectively. Let $n = (n_0 + n_1)$ be the number of haplotypes in the population with a non-missing call. We computed the mean number of pairwise differences for this SNP as follows. First, we computed the total number of pairs as $n_{pairs} = n * (n-1)/2$. Then, we computed the number of pairs that were the same, $n_{same} = (n_0 * (n_0 - 1)/2) + (n_1 * (n_1 - 1)/2)$, and the number of pairs that were different, $n_d = n_{pairs} - n_{same}$. Finally, we obtained the mean number of pairwise differences as $mpd = n_d/n_{pairs}$. To estimate nucleotide diversity $\pi$, we computed the sum of $mpd$ in a window of 20kbp centred on each SNP, and divided by the number of accessible bases, which produces the mean number of pairwise differences per base.

Regions containing high leverage points tended to be at the ends of chromosomes or within regions of high nucleotide diversity, where read mapping was problematic (see Figure S3.3). We identified potential outliers in all samples, and filtered out common outliers in at least 50 samples – 48,443 in total.

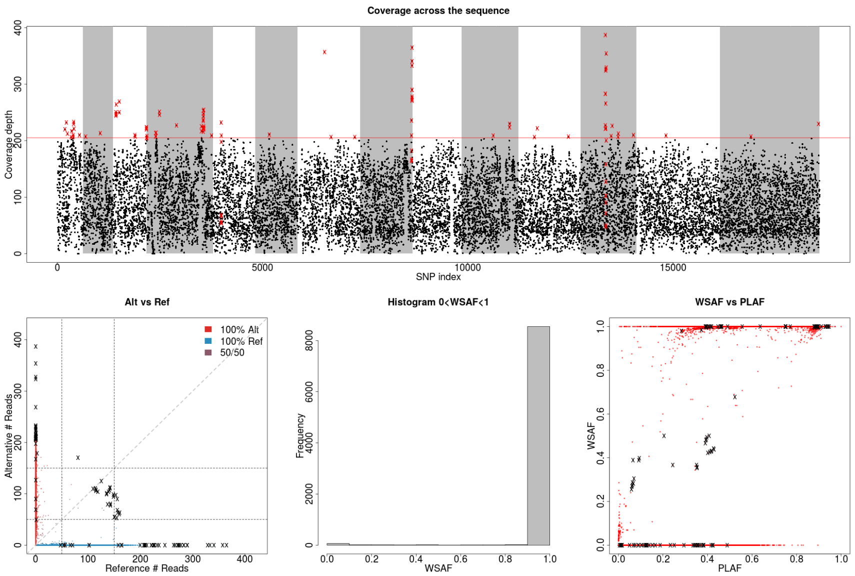


Figure S3.2: Identification of high leverage data points for filtering. (Top) Plot showing total allele counts across all markers for field isolate PG0415. We observe a small number of heterozygous sites with high coverage (shown as crosses on the bottom-left plot), which can potentially mislead our model to over-fit the data with additional strains (above the dotted line). We used a threshold of $\geq 99.5\%$ coverage to identify markers with high allele counts. Red crosses indicate markers that are filtered out. (Bottom-left) Scatter plot showing alternative against reference allele count. The marked black crosses refer to the outliers identified on the previous plot, which will cause the inference method to mistakenly identify the sample as being a mixed infection. (Bottom-middle) Histogram of allele frequency within sample. (Bottom-right) Allele frequency within sample (WSAF), compared against the population average (PLAF).

#### S3.3 Analysis preparation

To improve the accuracy and efficiency of the deconvolution process, we first split the data into groups based on genetic similarity. We computed genetic distance between two samples as follows:

$$d(x,y) = \sum_{l}^{L} \mathrm{WSAF}_{x,l} * (1 - \mathrm{WSAF}_{y,l}) + \mathrm{WSAF}_{x,l} * (1 - \mathrm{WSAF}_{y,l}) \tag{S3.7}$$

where $l$ represents an arbitrary locus, $L$ denotes the total number of loci, and $\mathrm{WSAF}_{s,l}$ indicates the non-reference within-sample allele frequency for sample $s$ at locus $l$. $\mathrm{WSAF}_{s,l}$ is then given by $\mathrm{WSAF}_{s,l} = \frac{a_{s,l}}{r_{s,l}+a_{s,l}}$ where $a_{s,l}$ is the number of read counts supporting the alternative allele in sample $s$ at locus $l$, and $r_{s,l}$ is the number of read counts supporting the reference allele in sample $s$ at locus $l$.

We found that samples from the same geographical region differentiated into clear groups. We used this initial grouping as the basis for defining the reference panels that assisted the deconvolution. The geographical groups arising from this analysis are listed below. In order to reduce computational time, we only used polymorphic sites at each population group:

1. Malawi, Congo, with 349,242 sites.
2. Ghana (Navrongo), with 508,606 sites.
3. Nigeria, Senegal, Mali, with 210,819 sites.
4. The Gambia, Guinea, Ghana (Kintampo), with 250,827 sites.

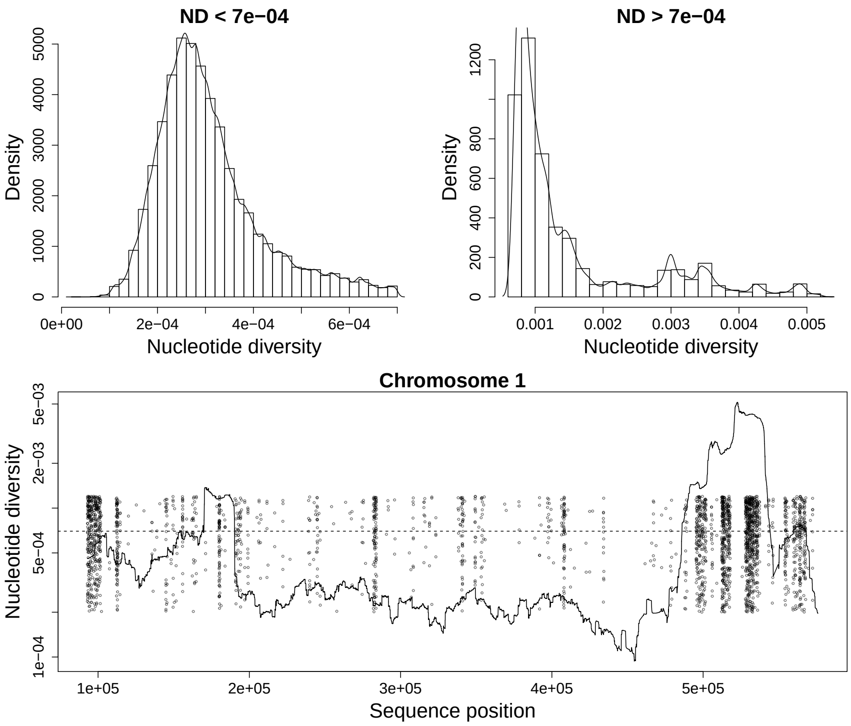


Figure S3.3: Nucleotide diversity for a sliding window size of 20,000 base pairs. (Top) Histograms showing the heavy tail of ND beyond 0.0007. (Bottom) Figure showing ND along *P. falciparum* chromosome 1. Scattered Points mark chromosome positions of poorly genotyped SNPs which we exclude from the deconvolution process. These points are jitterred to ease visualization.

5. Cambodia (Pursat), Cambodia (Pailin), Thailand (Sisakhet), with 44,317 sites.
6. Vietnam, Laos, Cambodia (Ratanakiri), Cambodia (Preah Vihear), with 88,410 sites.
7. Bangladesh, Myanmar, Thailand (Mae Sot), Thailand (Ranong), with 84,868 sites.

#### S3.4 Haplotype quality assessment

In this work, we also assessed the quality of the haplotypes inferred by `DEploidIBD`. Our goal was to establish to what degree our inferred haplotypes were statistically indistinguishable, given a suite of population genetics statistics, from the subset of clonal haplotypes that had the same geographical origin. Our assumption was that haplotypes found in mixed infections would have similar characteristics than those present in clonal samples. In our assessment, we found that the distribution of statistics for groups of deconvoluted haplotypes had extreme outliers and presented a higher variance when compared with the clonal population originating on the same region. We noticed that the painting process implemented by DEploid struggles when faced with challenging mixtures. For instance, mixed infections in which the co-existing strains have the same relative proportion (e.g. $k = 4$ with each strain having a proportion of 25%), or samples in which proportions are very unbalanced (e.g. k=2 with the marginal strain at 2%). This often results in an excess of alternative calls being assigned to one of the strains, which in turn provokes a deficit of diversity on the remaining haplotypes, that cannot be explained in terms of their genetic relationship to the reference genome used for mapping and assembly (3D7). We defined our quality metric as a $z$-score that approximates how much a deconvoluted haplotype deviates from the mean genetic diversity of the clonal population present in the same geographical area.

For each population, we computed the distribution of alternative calls observed within the subset of clonal samples ($k = 1$). Using this distribution as reference, we computed a $z$-score for each haplotype in the whole population following

$$z_i = \frac{a_i - \bar{a}_r}{\sigma_r},$$

where $a_i$ denotes the number of alternative calls in the haplotype $i$, and $\bar{a}_r$ and $\sigma_r$ are, respectively, the mean and standard deviation of observed alternative calls in the clonal set of samples from the population of origin. We only considere as suitable haplotypes with a $z$-score in the range $(-3, 3)$, thus discarding any strain that is three or more standard deviations away, in terms of alternative calls, from the mean observed for clonal samples. By using the set of clonal samples as the reference distribution, we approximated the number of alternative calls expected in a genome belonging to that population, which serves as a proxy for genetic diversity but is easier to compute. Supp. Figure S3.4 shows an example of this filtering process for the most problematic population in the dataset (Ghana). Supp. Table S3.3 lists the number of haplotypes discarded by population while Supp. Table S3.4 describes the number of haplotypes discarded by COI level. Statistical deconvolution of haplotypes in mixed infections remains a challenging problem and requires further research. Nonetheless, our quality metric can guide other researchers in the process of discarding haplotypes that are clearly artefactual

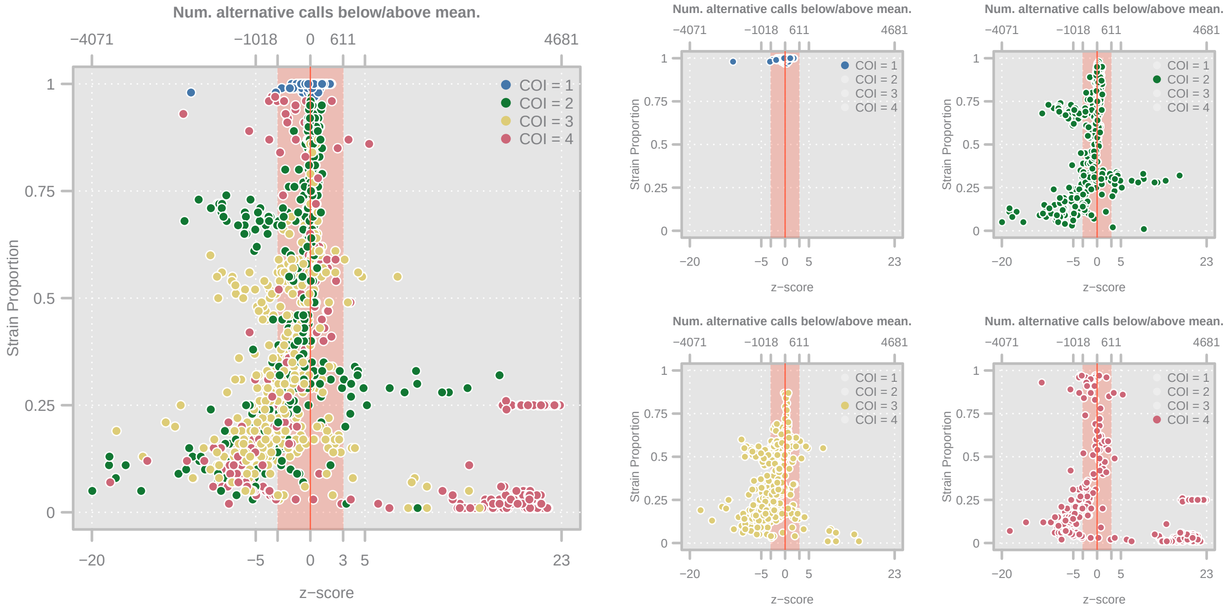


Figure S3.4: Diagnostic plots showing the distribution of haplotype quality ($z$-scores) for the Ghanian samples. Left. Scatterplot showing the relationship between haplotype $z$-score and strain proportion. The top axis shows the number of alternative calls below/above the mean of the subset of clonal samples that correspond to a given $z$-score. The vertical red line denotes a $z$-score of 0 whereas the red-shaded area indicate the haplotypes we retain Point colors show the COI level of the sample. Right. Four views of the same plot in which the samples have been highlighted according to their COI level.

#### S3.5 Combining clonal sample pairs for background IBD computation

We combined randomly selected clonal sample pairs to create artificial mixed infections, as a way to generate a background IBD distributions for each country. We assumed these artificial mixed infections mimic infections generated from two independent mosquito bites. In this setting, strain proportions are determined by their

| Country | Discarded | Retained | Fraction Discarded |
|---|---|---|---|
| Bangladesh | 25 | 69 | 0.27 |
| Cambodia | 108 | 697 | 0.13 |
| DR. of Congo | 62 | 155 | 0.29 |
| Ghana | 493 | 609 | 0.45 |
| Guinea | 79 | 88 | 0.47 |
| Laos | 28 | 110 | 0.20 |
| Malawi | 233 | 341 | 0.41 |
| Mali | 37 | 140 | 0.21 |
| Myanmar | 7 | 71 | 0.09 |
| Senegal | 2 | 167 | 0.01 |
| Thailand | 28 | 169 | 0.14 |
| The Gambia | 22 | 73 | 0.23 |
| Vietnam | 23 | 113 | 0.17 |
| **Total** | 1147 | 2802 | 0.29 |

Table S3.3: Number of haplotypes discarded and retained for each population in the Pf3k dataset.

| COI | Retained | Discarded | Fraction Discarded |
|---|---|---|---|
| 1 | 1331 | 34 | 0.02 |
| 2 | 669 | 291 | 0.30 |
| 3 | 583 | 533 | 0.48 |
| 4 | 219 | 289 | 0.57 |
| **Total** | 2802 | 1147 | |
| **Fraction** | 0.71 | 0.29 | |

Table S3.4: Number of haplotypes retained and discarded stratified by COI level.

median read depth, whereas sample coverage is obtained by accumulating the reference and alternative allele counts of two clonal samples. Similar to `DEploidIBD` deconvolution, SNPs with very high coverage resulting from this process caused high leverage in the model. Additionally, the sample sequence depth and skewness were heterogeneous due to different sample preparation and sequencing protocols. We reduced the `DEploidIBD` filtering threshold from 99.5% to 80%, and used low recombination probabilities to avoid false IBD breakpoint inference. We validated our method using lab crosses (Miles et al., 2016), and compared the IBD block detection using Li and Stephens (2003)'s painting with parental strains and the `DEploidIBD` algorithm (Figure S3.5).

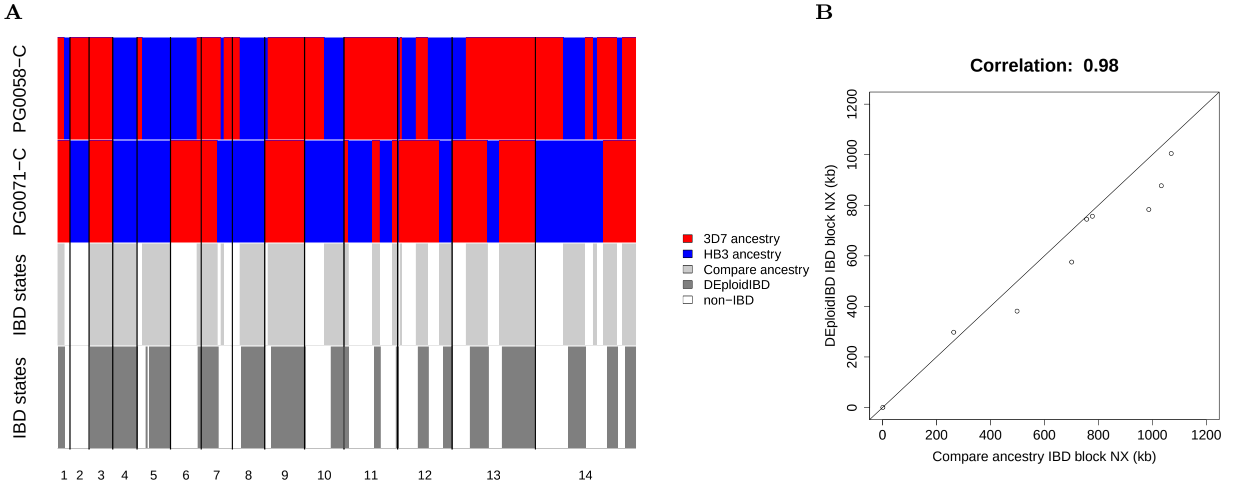


Figure S3.5: (A) Comparison of IBD block detection between DEploidIBD (top) and ancestral state inference from Li and Stephens (2003) (bottom), using artificial mixtures of lab crosses PG0071-C and PG0058-C (last tract). (B) Scatter plot of IBD segment Nx values extracted by comparing clonal sample ancestry (using DEploidIBD) on artificial mixtures.

### S4 Expected levels of IBD in *P.falciparum* mixed infections

The amount of IBD observed in a mixed infection is a function of the number of oocysts present in the biting mosquito. We will demonstrate this below.

First, let us briefly review the fundamentals of malaria meiosis. In our case, we imagine a mosquito bites a human host containing two distinct malaria strains. Call these strains $A$ and $B$. Some number of gametocytes of $A$ and $B$ are imbibed during the bite, differentiate into gametes, and undergo fertilization to produce zygotes (reviewed in Ghosh et al. (2000), Bennink et al. (2016)). Some fraction of these zygotes succeed in establishing themselves as oocysts on the mosquito midgut (Ghosh et al. (2000)). Three products of fertilization are possible, and thus the oocysts can be either: $A + A$ or $B + B$ (inbred oocysts, $n_{ii}$), or $A + B$ (outbred oocysts, $n_{ij}$). The oocyst state of a mosquito can be characterized by $(n_{ij}, n_{ii})$. Which strain is maternal and paternal may vary from oocyst to oocyst, but this is of no consequence here.

A $k = 2$ mixed infection is established when two distinct sporozoites, produced from the oocysts of this mosquito, infect a host. Each oocyst produces thousands of sporozoites (Beier et al. (1991)), of four types (discussed in McKenzie et al. (2001)), which pool in the mosquito salivary glands (Ghosh et al. (2000)). Imagine drawing a $k = 2$ mixed infection from a mosquito harbouring a single outbred oocyst ($n_{ij} = 1$). In such a mosquito there are two copies of each of the two strains (two sets of sister chromatids; $A$, $A$, $B$, $B$). Thus, ignoring recombination for the present, there are two pairs with an IBD fraction of 1 and, if our original strains are unrelated, the remainder of the $\binom{4}{2}$ pairs will have an IBD of 0. Thus a single $n_{ij}$ oocyst has an expected IBD of $E[\rho] = 2/\binom{4}{2} = 1/3$. We draw pairs without replacement because if sporozoites of only one type seed the infection, it will be $k = 1$. Importantly, neither recombination nor segregation change this result, as they only shuffle how the total identity is distributed between pairs, rather than create or destroy identity (identity is created by DNA replication and destroyed by mutation); the expectation is taken over all pairs and is thus unaffected.

Computing the expected IBD fraction for a mosquito possessing $n_{ij}$ outbred oocysts is an extension of the above. Again ignoring recombination, the expected IBD fraction $E[\rho|n_{ij}]$ is equal to the total number of pairs with an IBD of 1 (IBD pairs), over all possible pairs. In a mosquito with $n_{ij}$ oocysts, we have $2n_{ij}$ copies of each parental strain, thus we have $\binom{2n_{ij}}{2}$ IBD pairs for each parental strain, thus $2\binom{2n_{ij}}{2}$ IBD pairs total. Dividing this by the total number of pairs amongst $n_{ij}$ oocysts we have

$$E[\rho|n_{ij} > 0] = \frac{2\binom{2n_{ij}}{2}}{\binom{4n_{ij}}{2}} = \frac{2n_{ij}-1}{4n_{ij}-1}. \tag{S4.8}$$

The above yields 1/3 for $n_{ij} = 1$, approaching 1/2 as $n_{ij}$ grows. This result has been validated with `pf-meiosis` in Figure S4.6.

Including $n_{ii}$ oocysts is somewhat involved, as some pairs (selected without replacement) may be identical (thus yielding $k = 1$) or completely unrelated (yielding $k = 2$, but effectively without having undergone meiosis or producing any detectable recombination breakpoints between parental strains). We are interested in the expected IBD produced as a result of meiosis between parental strains, and thus for the moment we exclude these pairs. In practice, this means the mosquito must have at least one outbred oocyst, and at least one of the infecting sporozoites must be from an outbred oocyst.

The derivation is as above: first ignoring recombination and segregation, then enumerating all IBD pairs (pairs with IBD fraction of 1) and dividing by the total number of pairs to compute the expectation. Note that the additional IBD pairs possible between an outbred and inbred oocyst are given by the term $8n_{ij}n_{ii}$, and that you can no longer use all possible pairs drawn without replacement as the denominator, but must exclude the pairs described above.

$$\begin{aligned}E[\rho|n_{ij} > 0, n_{ii}] &= \frac{2\binom{2n_{ij}}{2} + 8n_{ij}n_{ii}}{2\binom{2n_{ij}}{2} + 16n_{ij}n_{ii} + 4n_{ij}^2} \\ &= \frac{2(n_{ij}+2n_{ii})-1}{4(n_{ij}+2n_{ii})-1}\end{aligned} \tag{S4.9}$$

Which is of a similar form to above, but increases to 1/2 quicker if more inbred oocysts are present. As before the equation is validated in Figure S4.7A.

The expression for $E[\rho|n_{ij} > 0, n_{ii}]$ can also be derived by recognizing that there are three *types* of pairs possible in a mosquito with a collection of $n_{ij}$ and $n_{ii}$ oocysts: (1) a pair can contain two strains from a single $n_{ij}$, ($n_{ij}^{o=1}$); (2) a pair can contain two strains from two different $n_{ij}$, ($n_{ij}^{o=2}$); or (3) a pair can contain one strain from an $n_{ij}$ oocyst and one from an $n_{ii}$ oocyst, $n_{ij,ii}^{o=2}$. Pair type (1) is unique to malaria and has an $E[\rho|n_{ij}^{o=1}] = 1/3$, as shown above; pair type (2) are standard siblings with $E[\rho|n_{ij}^{o=2}] = 1/2$; and pair type (3) represent a mother-daughter relationship, also with $E[\rho|n_{ij,ii}^{o=2}] = 1/2$. The full IBD fraction and IBD segment length distributions of these pairs were generated using `pf-meiosis` and can be seen in Figure S4.7B. We enumerate the number of each pair type given $n_{ij}$ and $n_{ii}$, weighted by their expectation, to derive $E[\rho|n_{ij} > 0, n_{ii}]$:

$$\begin{aligned}E[\rho|n_{ij} > 0, n_{ii}] &= \frac{n_{ij}^{o=1}E[f|n_{ij}^{o=1}] + n_{ij}^{o=2}E[f|n_{ij}^{o=2}] + n_{ij,ii}^{o=2}E[f|n_{ij,ii}^{o=2}]}{n_{ij}^{o=1} + n_{ij}^{o=2} + n_{ij,ii}^{o=2}} \\ &= \frac{n_{ij}\binom{4}{2}1/3 + 16\binom{n_{ij}}{2}1/2 + 16n_{ij}n_{ii}1/2}{n_{ij}\binom{4}{2} + 16\binom{n_{ij}}{2} + 16n_{ij}n_{ii}} \\ &= \frac{2(n_{ij}+2n_{ii})-1}{4(n_{ij}+2n_{ii})-1}\end{aligned} \tag{S4.10}$$

As above.

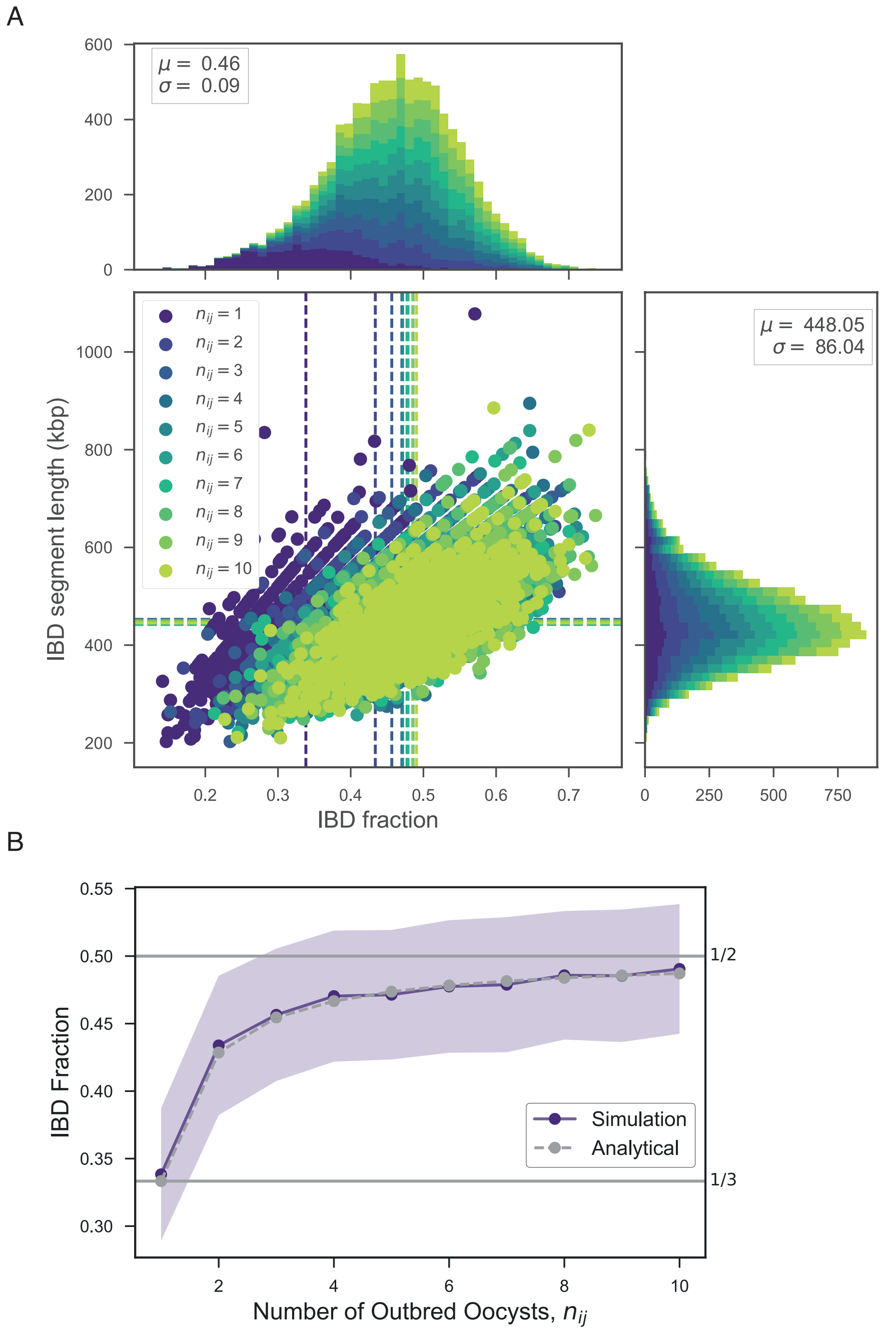


Figure S4.6: Exploring the relationship between number of outbred oocysts ($n_{ij}$) and IBD. (A) Joint IBD fraction and IBD segment length distributions for $k = 2$ mixed infections simulated from two unrelated strains and a fixed number of outbred oocysts $n_{ij}$, using `pf-meiosis`. Mean values for each distribution are indicated by same-color dashed lines. Each distribution is created from 1000 simulated mixed infections. (B) Validation of theoretical result given in text (S1.8). Line plot compares trend in expected IBD fraction with the number of outbred oocysts, $n_{ij}$, for infections simulated in panel A, and analytical expression S1.8.

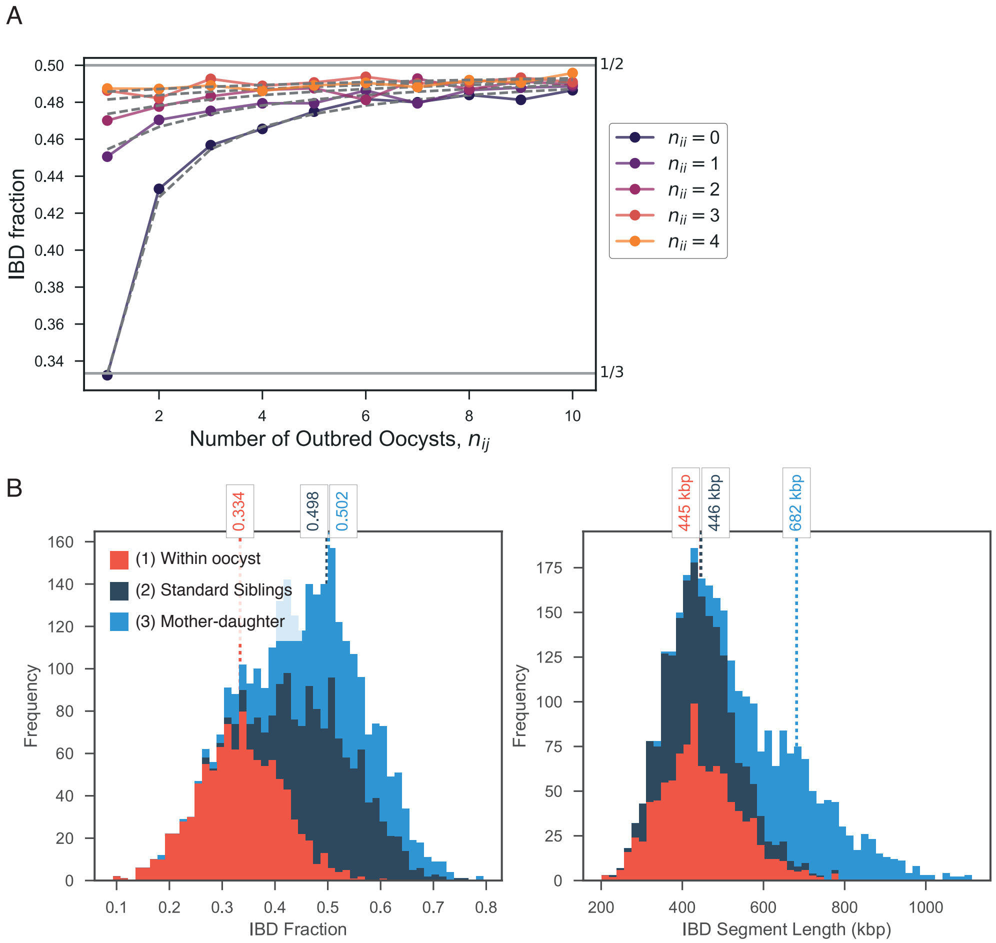


Figure S4.7: Exploring expected IBD allowing for outbred ($n_{ij}$) and inbred ($n_{ii}$) oocysts (A) Validation of expression for expected IBD fraction conditional on outbred $n_{ij}$ and inbred $n_{ii}$ oocysts (S1.9). Line plot compares trend in expected IBD fraction with varying number of outbred (x-axis, $n_{ij}$) and inbred (line color, $n_{ii}$) oocysts and the analytical expression S1.9 (grey dashed lines). (B) Using `pf-meiosis` to simulate $k = 2$ mixed infections generated from (1) two strains from the same outbred oocyst from ($n_{ij}^{o=1}$, 'Within oocyst'); (2) two strains different outbred oocysts($n_{ij}^{o=2}$, 'Standard Siblings'); (3) one strain from an outbred and one strain from an inbred oocyst ($n_{ij,ii}^{o=2}$, 'Mother-daughter').

### About the Pf3k Project

*The analyses in this paper are based on the Pf3k version 5 dataset, released in February 2016, comprising Illumina short read sequence data and SNP/indel calls on 2,512 field samples, 5 cultured parasite lines, 96 samples from genetic crosses, and 27 samples from artificial mixtures of cultured parasite lines.*

#### Background

The Pf3k Project was established in 2014 to develop high quality open access data resources for analysis of *Plasmodium falciparum* genome variation, based on standardised protocols for variant discovery and genotype calling, together with new tools for analysis of complex genetic variation in field samples (https://www.malariagen.net/projects/pf3k ).

Field samples from Bangladesh, Cambodia, Democratic Republic of Congo, Guinea, Laos, Malawi, Mali, Myanmar, Nigeria, Thailand and The Gambia were sequenced as part of the MalariaGEN *Plasmodium falciparum* Community Project (https://www.malariagen.net/projects/p-falciparum-community-project ). As these samples were collected by different MalariaGEN partner studies over a period of several years, and the sequence data have been made openly available, previous analyses of subsets of these samples can be found in publications by the MalariaGEN network and its partner studies (refs 1-12). Further details of individual partner studies are given below.

All sequencing data were produced by the Wellcome Sanger Institute (refs 1-12,15,16), with the exception of 137 samples from Senegal that were sequenced by the Broad Institute (ref 13,14). The sequence data on parents and progeny of genetic crosses have been previously published by Miles et al, 2016 (ref 15). The artificial mixtures of cultured parasite lines were contributed by Jason Wendler and Kirk Rockett at the Wellcome Trust Centre for Human Genetics and the Wellcome Sanger Institute. In another part of the Pf3k Project, PacBio long read sequence data were produced on 15 cultured isolates from different geographical locations, as reported by Otto et al, 2018 (ref 16).

Pf3k Project data can be openly accessed at https://www.malariagen.net/data and can be browsed using the interactive Panoptes web application (ref 17) at https://www.malariagen.net/apps/pf3k/ .

#### Members of the Pf3k Project

**Steering group**: Matt Berriman, Dominic Kwiatkowski, Gil McVean, Dan Neafsey

**Population genetic working group**: Jacob Almagro-Garcia, Roberto Amato, Zam Iqbal, Dominic Kwiatkowski, Sorina Maciuca, Gil McVean, John O'Brien, Joe Zhu.

**Technology benchmarking working group**: Roberto Amato, Dominic Kwiatkowski, Dan Neafsey, Richard Pearson, Jim Stalker

**Reference genome working group**: Matt Berriman, Chris Newbold, Thomas Otto

**Coordination**: Vikki Cornelius, Bronwyn MacInnis, Dawn Muddyman

**Partner study lead investigators**: Alfred Amambua-Ngwa, Lucas Amenga-Etego, Gordon Awandare, David Conway, Alistair Craig, Nicholas Day, Abdoulaye Djimde, Arjen Dondorp, Rick Fairhurst, Olivo Miotto, Nick White

#### MalariaGEN partner studies contributing to the Pf3k Project

Study 1001: Developing the Community Project with partners in Mali

- Description: https://www.malariagen.net/network/where-we-work/1001-developing-community-project-partners-mali
- Key contact: Abdoulaye Djimdé

Study 1006: Genome-wide analysis of genetic variation in The Gambia

- Description: https://www.malariagen.net/network/where-we-work/1006-genome-wide-analysis-genetic-variation-gambia
- Key contact: Alfred Amambua-Ngwa

Study 1017: Population genetics of natural populations in Northern Ghana

- Description: https://www.malariagen.net/network/where-we-work/1094-population-genetics-p-falciparum-parasites-northern-ghana
- Key contact: Lucas Amenga-Etego

Study 1022: Genome variation and selection in clinical isolates from rural Malawi

- Description: https://www.malariagen.net/network/where-we-work/1022-genome-variation-and-selection-clinical-isolates-rural-malawi
- Key contact: Alister Craig

Study 1026: Effects of transmission intensity on population structure and signatures of selection in Guinea

- Description: https://www.malariagen.net/network/where-we-work/1026-effects-transmission-intensity-population-structure-and-signatures
- Key contact: David Conway

Study 1044: Genomics of Plasmodium clearance and recrudescence rates in Cambodia

- Description: https://www.malariagen.net/network/where-we-work/1044-genomics-plasmodium-clearance-and-recrudescence-rates-cambodia
- Key contact: Thomas E. Wellems

Study 1052: Tracking Resistance to Artemisinin Collaboration (TRAC)

- Description: https://www.malariagen.net/network/where-we-work/1052-tracking-resistance-artemisinin-collaboration-trac
- Key contact: Elizabeth Ashley

Study 1083: Alternative molecular mechanisms for erythrocyte invasion by P. falciparum in Ghana

- Description: https://www.malariagen.net/network/where-we-work/1083-alternative-molecular-mechanisms-erythrocyte-invasion-p-falciparum-ghana
- Key contact: Gordon Awandare

Study 1094: Population genetics of P. falciparum parasites in Northern Ghana

- Description: https://www.malariagen.net/network/where-we-work/1094-population-genetics-p-falciparum-parasites-northern-ghana
- Key contact: Lucas Amenga-Etego
